# Larval antibiosis to cabbage stem flea beetle (*Psylliodes chrysocephala*) is absent within oilseed rape (*Brassica napus*)

**DOI:** 10.64898/2026.01.22.694425

**Authors:** Ryan E. Brock, Clara Courtney, Steven Penfield, Rachel Wells

## Abstract

**BACKGROUND:** Insect pests present a global threat to crops, with plant resistance representing a key breeding goal. The cabbage stem flea beetle (*Psylliodes chrysocephala*; CSFB) is a key pest of oilseed rape (*Brassica napus*; OSR) within Europe; however, CSFB resistance is yet to be found within *B. napus*. To address this, we examine CSFB larval development over time, explore antibiosis across a genetically diverse *B. napus* panel, and test whether larvae can develop within model Brassicaceae species (*Brassica rapa* and *Arabidopsis thaliana*).

**RESULTS:** CSFB larvae completed development from four-weeks post-infestation, undergoing a 20-fold size increase, with larval recovery after two weeks allowing semi-high throughput resistance phenotyping. Applying this method to 98 Brassicaceae genotypes (97 *B. napus* and a single *Sinapis alba*), we found weak evidence for genotype effects on larval survival. However, phenotype validation with ‘resistant’ and ‘susceptible’ *B. napus* genotypes showed no differences in larval survival or adult emergence. Larval antibiosis was consistently observed in *S. alba*. Finally, we showed that model *B. rapa* and *A. thaliana* genotypes represent suitable hosts for CSFB, with larvae increasing 8-10× in size after two weeks.

**CONCLUSION:** CSFB larval antibiosis appears absent within *B. napus*, possibly due to bottlenecks experienced during domestication. However, larval antibiosis is present within *S. alba*, and future work should study the basis of this resistance. Further, CSFB larval screening in Brassicaceae model species presents an opportunity to explore CSFB resistance genetics, informing breeding progress for insect resistance in *B. napus*.

## 1. Introduction

Insect herbivory is one of the most common trophic interactions on earth and represents a major selection pressure for plant defence traits.^1–3^ In food crops, selective breeding for high yield, quality traits, and uniform growth has reduced crop genetic diversity and altered plant-m insect interactions,^4–6^ such that modern crops are generally more insect-susceptible than their wild relatives.^7,8^ In addition, agricultural ecosystems often comprise monocultures of crop genotypes, further increasing insect susceptibility.^9^ Insect pests therefore represent a global threat to food production, causing assumed yield losses of up to 40% a year.^10,11^

Pesticide application broadly limits pest impacts on crop yield and enhances agricultural productivity^12,13^ (although see Perrot *et al.*^14^). However, the adverse environmental effects of pesticide use^15,16^ coupled with the evolution of pesticide-resistant insect populations^17^ mean more sustainable pest management approaches are required. Host plant resistance, defined as plant traits influencing insect-associated damage,^18^ offers a promising strategy for pest management.^19–21^ Plant resistance to insect herbivory can manifest through three non-exclusive pathways: (1) antixenosis, in which insects are deterred from host plant colonisation; (2) antibiosis, in which insects suffer reduced fitness on host plants; and (3) tolerance, in which plants compensate for insect damage through increased growth or reproduction.^18,21,22^ A crucial first step in identifying host plant resistance requires phenotyping for resistance variation across plant species or genotypes, before using genetic mapping approaches to associate genes and traits with resistance.^20,21^ However, the identification of insect resistance genes has proven challenging due to the polygenic nature of resistance and labour-intensive phenotyping methodologies.^21,23,24^

Oilseed rape (*Brassica napus*; OSR), an allopolyploid derived from the hybridisation of *Brassica rapa* and *Brassica oleracea*,^25^ represents the second-largest oilseed crop by area grown globally and a key break crop in rotation systems.^26,27^ An important breakthrough in OSR commercial success was the production of double-low genotypes,^25,28,29^ plants yielding seeds with low levels of both erucic acid, a fatty acid with potential cardiotoxic effects,^30^ and glucosinolates, a group of secondary metabolites involved in plant defence.^31^ As a result, modern OSR genotypes are severely bottlenecked and express low levels of glucosinolates within tissues,^25,32–35^ compromising disease and pest resistance. OSR is therefore susceptible to a range of pathogens and insects,^36,37^ with insect resistance now representing a major objective for breeders.^23,38^

Within Europe, the cabbage stem flea beetle (*Psylliodes chrysocephala*; CSFB), a Brassicaceae specialist,^39^ represents a key pest of OSR.^37^ The CSFB life cycle is closely linked to the cropping cycle of winter OSR, with distinct damage symptoms associated with different life stages.^40^ Adult CSFBs feed on emerging seedlings during early autumn, impacting crop establishment and reducing vigour.^40^ Following adult reproduction, CSFB larvae mine OSR stems and petioles during autumn and winter, passing through three larval instars before leaving the plant to pupate in spring.^40^ CSFB larvae are the more economically damaging life-stage, with mining leading to stunted growth, frost and pathogen susceptibility, and plant mortality, causing severe yield penalties.^41–44^

CSFB control within OSR is heavily pesticide reliant. Previously, management involved the use of neonicotinoid seed treatments that provided systemic protection throughout the growth cycle.^45,46^ However, concerns over the environmental effects of neonicotinoids have led to usage restrictions within the European Union since 2013.^47,48^ In response, growers have become increasingly reliant on pyrethroid insecticides,^45,46^ driving the evolution of pyrethroid resistance within European CSFB populations.^49–51^ Limited control options have therefore led to increased CSFB abundance and subsequent reductions in OSR growth.^46,52,53^ In the UK, a ten-fold increase in CSFB larval loads^52^ has been accompanied by a 67% reduction in OSR land use (from 621,000ha in 2014 to 204,000ha in 2025)^54^ following the neonicotinoid ban. Given policies aimed at reducing pesticide use,^55^ the development of CSFB-resistant OSR genotypes represents a key target for the European OSR industry.^23,38^

Reported attempts to phenotype for CSFB resistance within Brassicaceae have been extremely limited to date, with just a single study comparing larval resistance between *B. napus* and four relatives (*B. rapa*, *B. oleracea*, *Sinapis alba*, and *Raphanus sativa*).^56^ This is mirrored for other insect pests of *B. napus* (reviewed in Hervé^23^ and Obermeier^38^), reflecting the challenging nature of exploring insect resistance. Despite phenotyping relatively small Brassicaceae collections, these studies have highlighted a lack of insect resistance across *B. napus* genotypes, partial resistance in some diploid relatives (*B. rapa* and *B. oleracea*) and resynthesised *B. napus* genotypes, and resistance and/or tolerance in more distantly related Brassicaceae species.^23,38^ CSFB larval antibiosis has been shown within *S. alba*, with larvae experiencing lower survival and slower development, while such effects were lacking across four tested *B. napus* genotypes.^56^ However, whether CSFB larval antibiosis exists across a more genetically diverse *B. napus* population remains to be seen.

In the present study, we aimed to explore CSFB larval antibiosis (defined as reduced survival and development) across a genetically diverse *B. napus* population, with the goal of identifying resistant genotypes that may provide trait and gene discovery for future breeding efforts. We outline a method for CSFB larval screening^57^ and investigate larval development under laboratory conditions, with our results informing a semi-high throughput approach for phenotyping CSFB larval antibiosis across 98 Brassicaceae genotypes (97 *B. napus* and a single *S. alba*). Importantly, this represents the largest *B. napus* collection phenotyped for insect resistance to date. Given the phenotypic diversity within this collection,^58–60^ we hypothesised that larval antibiosis would exist within some *B. napus* genotypes, providing a foundation for association mapping of genes involved in CSFB resistance. Finally, to broaden the genetic resources available for CSFB-Brassicaceae interactions, we applied CSFB larval screening to two model Brassicaceae species, *B. rapa* and *Arabidopsis thaliana*.

## 2. Materials and Methods

### 2.1. Cabbage stem flea beetle rearing

We established laboratory CSFB populations by rearing larvae from OSR (KWS Barbados) collected at the John Innes Centre Field Station (Bawburgh, Norfolk, UK) in the winters of 2021/2022 and 2022/23. Larvae were recovered from OSR plants using the Berlese method^61^ and placed onto pak choi (*B. rapa chinensis* Hanakan, Suttons Seeds, Paignton, UK) to complete development. On emergence, adult beetles were kept in populations of up to 70 individuals inside ventilated boxes (L17×W12×H6cm) lined with moist tissue and fed Chinese cabbage (*B. rapa pekinensis* Hilton, Suttons Seeds) *ad libitum*. Fresh cabbage was provided every 7-12 days, during which dead adults were removed and eggs were collected. Collected eggs were placed on pak choi to undergo development, with newly emerged adults used to setup new populations, thereby completing the CSFB lifecycle under laboratory conditions. All rearing steps were carried out under constant conditions (22±1°C, ∼60% RH, L16:D8). Captive CSFB populations would have cycled through approximately 3-9 generations when the below experiments were carried out, with no further addition of field-collected individuals since winter 2022/23.

### 2.2. Larval development time course

To understand CSFB larval development under laboratory conditions, we conducted a time course experiment across four timepoints (1-4 weeks after infestation) in a single *B. napus* genotype (Apex-93_5×Ginyou_3 DH; Warwick Gene Bank accession = 12841-mc-6). Seeds were sown into Levington Advance F2 compost (ICL, Amsterdam, NL) and ten days post-sowing, 20 seedlings were individually repotted into 9cm pots containing JIC Cereal Mix (65% peat, 25% loam, 10% grit, 3kg/m^3^ dolomitic limestone, 1.3kg/m^3^ PG Mix, 3kg/m^3^ osmocote exact), assigned to a timepoint, and randomised (**Figure S1**). Plants were grown under constant conditions (18±1°C, ∼60% RH, L10:D14) and watered regularly. The same growth procedure and conditions were used for all other experiments conducted.

Larval infestation began four-weeks post-sowing, when plants possessed 5-6 true leaves (mean = 5.3). Infestation was based on Döring & Ulber^56^ and Coston *et al.*^43^, with 12, <24 hour-old CSFB larvae added to each plant over a four-day period (i.e. three larvae per day),^57^ reflecting field-realistic UK infestation levels expected to lower harvest yield.^43,52,61^ Adding larvae across four-days allows sufficient larvae to hatch, is field-relevant (whereby larvae enter plants across multiple months),^42^ and ensures plants are not overly stressed from multiple larvae entering concurrently.

To stimulate hatching, approximately 1000 CSFB eggs were placed at constant conditions (22±1°C, ∼60% RH, L16:D8) 1-4 days prior to infestation. Each infestation day, three <24 hour-old larvae were transferred to the base of each plant (**Figure 1A**) and observed for 10 seconds to ensure movement. Any larvae that did not move were removed and replaced with a new larva. Each plant was sealed within a microperforated bag (L70×W38cm), ensuring that larvae could not move between plants. To account for circadian effects on plant defence compounds,^62^ larval infestation was carried out at the same time every day, with plants infested in a different order each day. On the final infestation day, plants were checked for successful larval penetration, defined as entry holes and mines visible on the stem or petioles (**Figure 1A**) and/or visible larvae under the plant epidermis.

**Figure 1.**
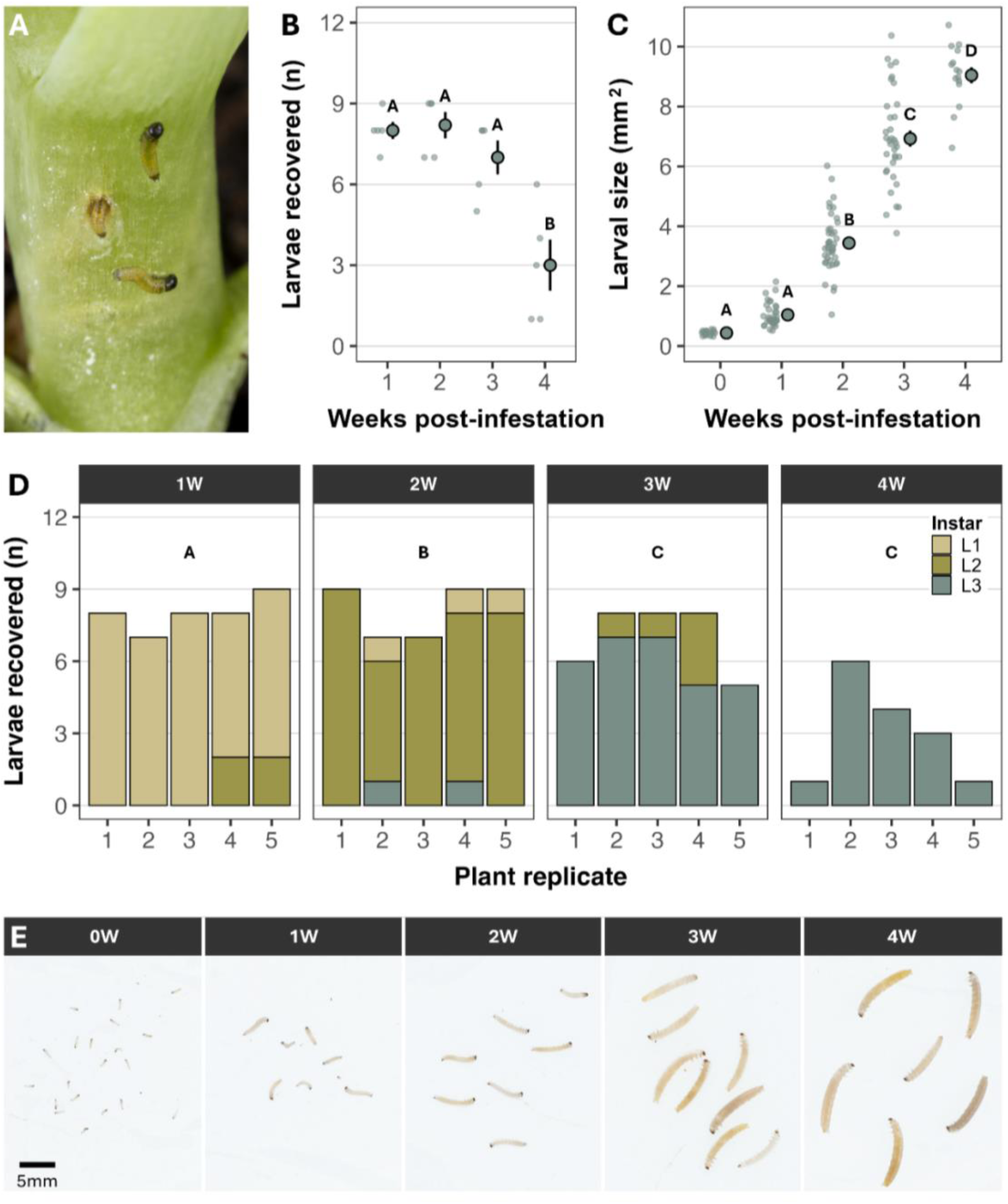
Cabbage stem flea beetle (*Psylliodes chrysocephala*) larval development within *Brassica napus* plants under laboratory conditions. **(A)** Infestation of *B. napus* with <24 hour-old CSFB larvae. **(B)** Live larval recovery from infested plants at each timepoint (1-4W; *n* = 5 plants per timepoint). **(C)** Larval size (mm^2^) at each timepoint. Week 0 represents the size of <24 hour-old larvae. For plots **B-C**, large points with whiskers represent the mean ± standard error, left jittered points represent the raw data, and letters represent significant differences (*p* < 0.05) between timepoints. **(D)** Larval instar proportions within each plant across timepoints. Light green bars = first instar (L1); mid-green bars = second instar (L2); dark green bars = third instar (L3). Letters within plots represent significant differences in instar composition (*p* < 0.05) between timepoints. **(E)** Representative photographs of larvae across timepoints.

Five infested plants were harvested and weighed at each timepoint, with the leaves, petioles and stem dissected to recover larvae. Location and instar of all live larvae within each plant were recorded, with recovered larvae stored in 70% ethanol until imaging. Dead larvae found during dissection were also recorded. Twenty <24 hour-old larvae were also collected for imaging. All collected larvae were photographed (**Figure S2A-B**), before ImageJ v1.52a^63^ was used to threshold the image and measure larval size (in mm^2^) with the ‘Analyze Particles’ function (**Figure S2C-D**). Occasionally, larvae were damaged during dissection and unavailable for measurement. Of the 131 live larvae recovered during dissections, 126 were measured.

### 2.3. Screening a genetically diverse *B. napus* population for larval antibiosis

To explore variation in CSFB larval antibiosis in *B. napus*, we conducted larval screening across 96 genetically fixed genotypes, comprising spring, winter and semi-winter OSR, swede and kale morphotypes, from the Renewable Industrial Products from Rapeseed^58^ panel (**Table S1**). Genotypes were randomised across a temporal 12-block alpha design (plots per block (*k*) = 24, replicates per genotype (*r*) = 3, blocks per replicate (*s*) = 4) and grown across two CER shelves, with a new block beginning every 1-9 weeks (**Table S2**; **Figure S3**). Each block also contained a negative *B. napus* control (KWS Campus) and a positive *Sinapis alba* control (Elsoms G1), with CSFB larval antibiosis having been previously demonstrated in *S. alba*.^56^ In a single block, two genotypes died before infestation and were therefore regrown in later blocks. Hence, each block contained 24-27 plants, giving a total of three replicates for all 96 *B. napus* genotypes and 12 replicates for both control genotypes.

Larval infestation^57^ was performed at four-weeks post-sowing, when plants possessed 1-9 true leaves (mean = 4.32). Our larval development experiment showed highest larval recovery two-weeks post-infestation, with larvae increasing eight-fold in size (**Figure 1B-C**). Hence, two-weeks post-infestation, all plants were harvested and weighed, with larvae recovered using the Berlese method^61^ at 22±1°C for up to 17 days. Of the 2,229 larvae recovered, 2,201 were imaged and measured as before. Larval antibiosis was defined from both the number of larvae recovered (i.e. larval survival) and the size of recovered larvae (i.e. larval development).

### 2.4. Phenotype validation

We identified potential variation in CSFB larval antibiosis across *B. napus* genotypes (**Figure 2A**). Therefore, to confirm larval antibiosis variation, we selected six genotypes (three ‘resistant’ – 33, 61 and 67 – and three ‘susceptible’ – 14, 84 and 89) for phenotype validation, along with the two control genotypes used previously. A total of nine replicates per genotype were phenotyped across three temporal blocks (i.e. three genotype replicates per block, giving 24 plants per block and 72 plants overall). Plants were grown in randomised blocks across two CER shelves, with a new block starting every 1-3 weeks (**Figure S4**). Larval infestation^57^ was performed three-weeks post-sowing, when plants possessed 2-6 true leaves (mean = 3.88). Two-weeks post-infestation, plants were harvested, weighed, and larvae were collected using the Berlese method,^61^ with all recovered larvae (*n* = 561) imaged and measured.

**Figure 2.**
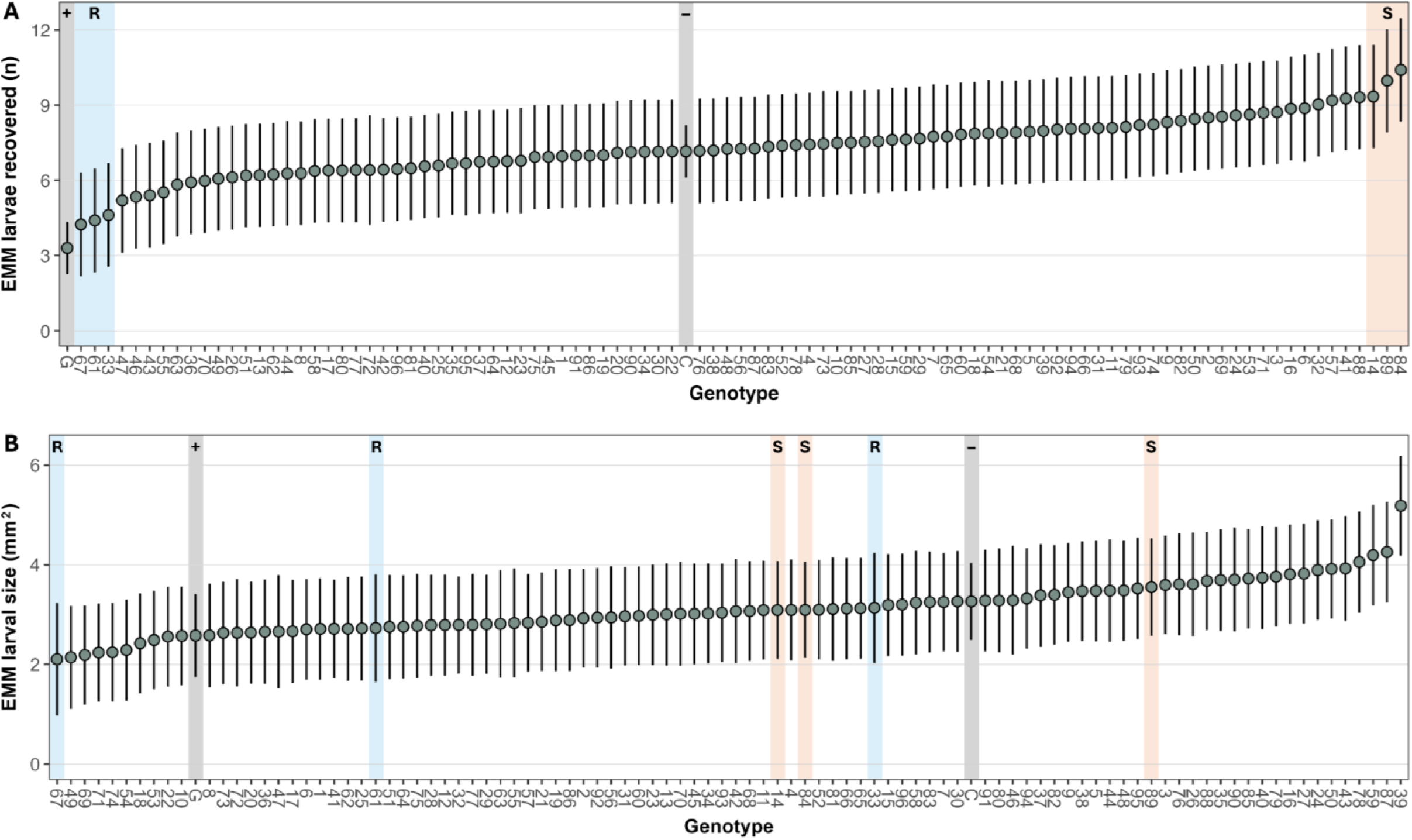
Per genotype estimated marginal mean (EMM) ± estimated 95% confidence intervals for cabbage stem flea beetle (*Psylliodes chrysocephala*) larval survival **(A)** and larval size (mm^2^) **(B)** across 98 Brassicaceae genotypes (97 *Brassica napus* and a single *Sinapis alba*). Larval survival is defined as the number of larvae recovered from a plant two-weeks post-infestation with 12 larvae. Genotypes are arranged from lowest to highest EMM. Blue and peach highlighted bars represent genotypes identified as resistant (R) and susceptible (S), respectively. Grey highlighted bars represent the positive (G = *S. alba* Elsoms G1) and negative (C = *B. napus* KWS Campus) controls.

To link larval recovery to adult development, we also explored adult emergence across the selected genotypes. Patterns of larval recovery between the control genotypes were similar across the *B. napus* population screen and phenotype validation experiment (**Figure S5**). Therefore, to increase throughput, the adult emergence experiment only contained the six *B. napus* genotypes of interest. A total of 10 replicates for each genotype were phenotyped across two temporal blocks (i.e. five genotype replicates per block, giving 30 plants per block and 60 plants overall). Plants were grown in randomised blocks across two CER shelves, with a new block starting each week (**Figure S6**). Larval infestation^57^ was carried out four-weeks post-sowing, when plants possessed 3-6 true leaves (mean = 4.25), and larvae were left to develop until adult emergence.

Based on larval development results (**Figure 1B-D**), we stopped watering plants at six-weeks post-infestation (since all larvae should have left the plants to pupate) and began checking for adult emergence every 2-7 days. Seven-weeks post-infestation (when adults were first observed), plant vegetation was disposed, allowing easier collection of adults. At each check, collected adults were counted and removed using an insect aspirator, with the experiment terminated when no new adults had emerged for two weeks. Hence, this experiment provided two extra measures of larval antibiosis: total adult emergence (i.e. the number of larvae that successfully completed development) and time to adult emergence (i.e. larval development time). One replicate of genotype 61 died between infestation and adult collection and so was excluded from analyses.

A full protocol for CSFB larval antibiosis phenotyping can be found in Brock & Wells.^57^

### 2.5. Larval screening in Brassicaceae model species

*Brassica napus* relatives may offer novel insights into CSFB resistance by providing tractable genetic resources.^23,56^ Hence, we assessed whether CSFB larval screening worked in reference *B. rapa* and *A. thaliana* genotypes.

*Brassica rapa* larval screening was carried out using the R-o-18 genotype, a rapid-cycling yellow mustard (*B. rapa trilocularis*) with genetic resources that include a high-quality genome and sequenced TILLING (targeting induced local lesions in genome) population.^64^ Plants were grown as per *B. napus*, with a total of 15 plants grown for larval screening. Infestation^57^ was carried out at three-weeks post-sowing (**Figure 4A**) when plants possessed 4-5 true leaves (mean = 4.20). Post-infestation, a single plant died, leaving 14 plants for larval screening. Two-weeks post-infestation, plants were harvested and larvae were recovered as before.^61^

*Arabidopsis thaliana* larval screening was carried out using the Ws-0 genotype. Seeds were sown into Levington Advance F2 and stratified at 4°C for one week, before being transferred to the same growth conditions used for *B. napus*. Three-weeks post-stratification, 15 *A. thaliana* plants were repotted into 5cm pots, and left to grow for a further week. Larval infestation was carried out four-weeks post-stratification, when rosette stage plants possessed 11-18 true leaves (mean = 14.13). Approximately 100 CSFB eggs were setup two days before larval infestation. Plants were infested by introducing a single <24 hour-old larva to the centre of the rosette (**Figure 4D**), with the introduced larva observed for 10 seconds to ensure movement. Plants were enclosed within sealed perforated bags (L30×W15cm), ensuring larvae could not move between plants. One-week post-infestation, plants were checked for successful larval infestation, defined as visible larval entry holes and/or mines on the petioles (**Figure 4E**). Two-weeks post-infestation, plants were harvested and larvae were collected as before.^61^

Recovered larvae (*B. rapa n* = 137; *A. thaliana n* = 9) and <24 hour-old larvae (*B. rapa n* = 12; *A. thaliana n* = 15) from both experiments were imaged and measured as before.

### 2.5. Statistical analyses

Analyses were conducted using R v4.5.2 in RStudio v2026.01.0.^65,66^ Linear models and linear mixed models were fitted using the ‘stats’ and ‘lmerTest’ packages,^67^ respectively. All plots were created with ‘ggplot2’.^68^ Linear model effects were assessed using the *Anova()* function (type II) from the ‘car’ package.^69^ Linear mixed model fixed effects were assessed using the *anova()* function (type II) from the ‘lmerTest’ package^67^, allowing *p*-value calculation via Kenward-Rogers approximation. Maximal models comprising all explanatory effects were fitted before non-significant effects (*p* > 0.05) were dropped in a stepwise manner to produce final models (**Table S3**). AIC-based model selection and assumption checks were performed using the *compare_performance()* and *check_model()* functions from the ‘performance’ package^70^. Estimated marginal mean (EMM) calculations and Tukey post hoc comparisons were performed using the ‘emmeans’ package^71^, with a significance threshold of *p* < 0.05. All R packages used for analyses are given in **Table S4**.

To analyse larval development over time, we fitted live larval recovery, larval size, and instar proportion as a function of timepoint. Instar size differences were analysed by fitting larval size as a function of instar. To analyse larval antibiosis across all Brassicaceae genotypes plus those selected for phenotype validation, we fitted larval recovery and larval size as a function of genotype. Development success during the adult emergence experiment was analysed by fitting total adult emergence, time (weeks) to first adult emergence, and mean adult emergence time as a function of genotype. Finally, to analyse larval development within *B. rapa* and *A. thaliana*, we fitted larval size as a function of status (day-old vs. recovered). Where necessary, models contained additional terms accounting for experimental design, larval nesting within plants, or confounding factors such as plant weight (see **Supplementary Methods** for full details).

Unless otherwise stated, values in the results section represent the raw mean ± standard error.

## 3. Results

### 3.1. Larval development occurs across four weeks under laboratory conditions

All infested plants (*n* = 20) showed CSFB larval herbivory symptoms. Live larval recovery was significantly influenced by plant weight (g) at dissection (F_(1, 15)_ = 5.16, *p* = 0.04) and timepoint (F_(3, 15)_ = 17.07, *p* < 0.001), with consistent larval recovery for the first three weeks (1W: 8.00±0.32; 2W: 8.20±0.49; 3W: 7.00±0.63) before decreasing to 3.00±0.95 at four-weeks post-infestation (**Figure 1B**). The same pattern was observed when including dead larvae (**Figure S7**). Larvae grew significantly over time (F_(4, 12)_ = 134.65, *p* < 0.001), undergoing a ∼20.5× size increase from 0.44±0.02mm^2^ at hatching to 9.05±0.27mm^2^ at four weeks (**Figure 1C & 1E**). Instar composition also changed significantly over time (F_(3, 16)_ = 368.51, *p* < 0.001; **Figure 1D**) and larval size significantly increased with instar (F_(2, 118)_ = 63.19, *p* < 0.001; **Figure S8**). These results suggest that larvae complete development and begin to pupate from four-weeks post-infestation under our laboratory conditions.

Plant dissections revealed changes in larval location over time, with larvae distributed across the stem, cotyledons, and petioles at one-week post-infestation, before feeding almost exclusively within petioles from 2-4 weeks (**Figure S9**). Petiole sharing was frequently observed, with 42.47% of infested petioles (31/71) containing two or more larvae (**Figure S10**).

### 3.2. Potential larval antibiosis within a genetically diverse *B. napus* population

All plants (*n* = 312) showed larval herbivory symptoms. Along with plant weight (g) at evacuation (F_(1, 144)_ = 6.72, *p* = 0.01), larval survival was significantly influenced by genotype (F_(97, 194)_ = 1.83, *p* < 0.001; **Table 1**), with lowest survival in *S. alba* G1 and highest survival in *B. napus* 84 (**Figure 2A**; **Figure S11A**). Importantly, larval survival was significantly lower (t_(1, 202)_ = −5.33, *p* < 0.001) in the *S. alba* positive control compared to the negative control (**Figure S12A**; **Figure S13**), demonstrating expected differences in CSFB larval survival between *S. alba* and *B. napus*.^56^ Excluding *S. alba*, EMM (95% CI) larval survival ranged from 4.24 (2.18-6.31) to 10.40 (8.34-12.47) across the 97 *B. napus* genotypes (**Figure 2A**), suggesting that genetic variation for CSFB larval antibiosis may exist within *B. napus*. Using our larval survival distribution, we assigned three genotypes as resistant (EMM (95% CI) larval survival: **67** = 4.24 (2.18-6.31); **61** = 4.40 (2.33-6.47); **33** = 4.62 (2.55-6.68)) and three as susceptible (EMM (95% CI) larval survival: **14** = 9.35 (7.28-11.41); **89** = 9.97 (7.91-12.03); **84** = 10.40 (8.34-12.47)). Importantly, larval survival was not significantly different between the three resistant genotypes and *S. alba*, but larval survival was significantly lower in the three resistant genotypes compared to the three susceptible genotypes (**Figure S13**).

**Table 1.** Fixed effect summaries for cabbage stem flea beetle (*Psylliodes chrysocephala*) larval survival and larval size across the 98 Brassicaceae plant genotypes (97 *Brassica napus* and a single *Sinapis alba*) screened for larval antibiosis.

| Response | Fixed effect | F | Num. df | Den. df | *p* |
| --- | --- | --- | --- | --- | --- |
| Larval survival | Genotype | 1.83 | 97 | 194 | **<0.001** |
|  | Plant weight | 6.72 | 1 | 144 | **0.01** |
|  | Replicate | 1.15 | 2 | 6 | 0.37 |
| Larval size | Genotype | 1.73 | 97 | 170 | **<0.001** |
|  | Replicate | 2.51 | 2 | 8 | 0.14 |

Num. df = numerical degrees of freedom; Den. df = Kenward-Roger estimated denominator degrees of freedom.

Larval size (mm^2^) was not influenced by plant weight or larval density within plants but did vary significantly by genotype (F_(97, 170)_ = 1.73, *p* < 0.001; **Table 1**), with smallest larvae in *B. napus* 67 and largest larvae in *B. napus* 39 (**Figure 2B**; **Figure S11B**). There was no significant difference in larval size between control genotypes (**Figure S12B**) nor between any of the resistant and susceptible genotypes identified by larval survival (**Figure S14**). Using per genotype raw means, we found a weak correlation between larval survival and larval size (F_(1, 96)_ = 4.11, *p* = 0.05; **Figure S15A**), suggesting that larvae may develop faster within genotypes in which survival is higher. However, this correlation was not present when using per genotype EMMs (F_(1, 96)_ = 4.11, *p* = 0.09; **Figure S15B**). Per genotype raw and estimated means for larval survival and size can be found in **Tables S5-S8**.

### 3.3. Phenotype validation did not confirm the presence of antibiosis

All plants in the phenotype validation experiment (*n* = 72) showed larval herbivory symptoms. Larval survival was again influenced by both plant weight (g) at evacuation (F_(1, 58)_ = 6.02, *p* = 0.02) and genotype (F_(7, 59)_ = 17.77, *p* < 0.001; **Table 2**), with lowest survival in *S. alba* G1 and highest survival in *B. napus* 84 (**Figure 3A**). However, no significant differences in larval survival were found across any of the initially identified resistant and susceptible genotypes (**Figure 3A**), with the plant genotype effect driven entirely by lower larval survival within *S. alba* compared to *B. napus*.

**Figure 3.**
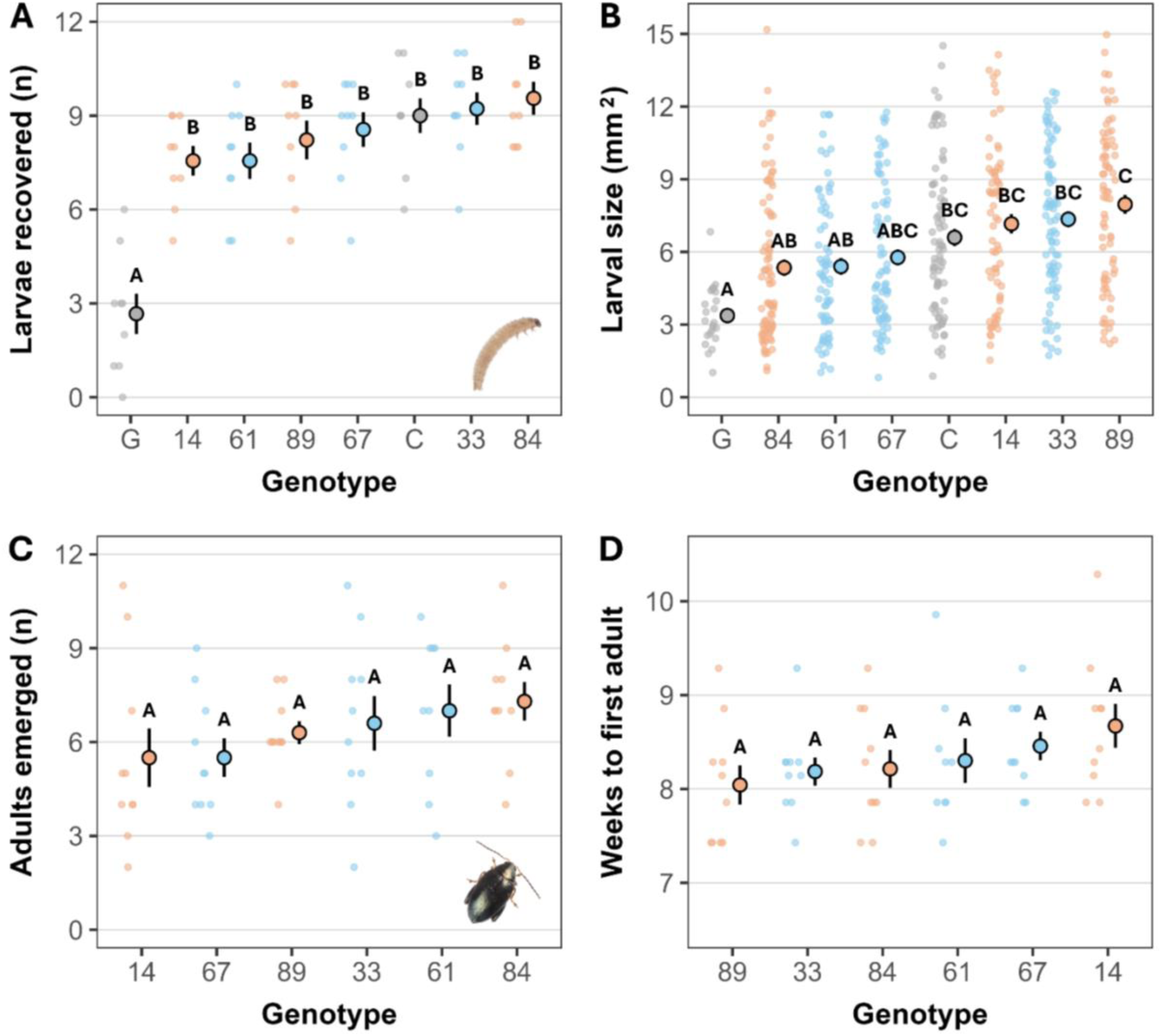
Cabbage stem flea beetle (*Psylliodes chrysocephala*) larval survival **(A)**, larval size **(B)**, and adult emergence **(C-D)** across eight selected Brassicaceae plant genotypes (seven *Brassica napus* and a single *Sinapis alba*). **(A)** Larval survival, defined as the number of larvae recovered from plants two-weeks post-infestation with 12 larvae, across eight Brassicaceae genotypes. **(B)** Larval size (mm^2^) across eight Brassicaceae genotypes. **(C)** Total adult emergence across six *B. napus* genotypes. **(D)** Time (weeks) to first adult beetle emergence across six *B. napus* genotypes. Genotypes are ordered from lowest to highest mean. Blue and peach points represent genotypes from the Brassicaceae population larval antibiosis screen that were initially identified as resistant and susceptible, respectively. Grey points represent the positive (G = *S. alba* Elsoms G1) and negative (C = *B. napus* KWS Campus) controls. Large points with whiskers represent the mean ± standard error, left jittered points represent the raw data, and letters represent significant differences (*p* < 0.05) between genotypes.

**Table 2.** Fixed effect summaries for cabbage stem flea beetle (*Psylliodes chrysocephala*) larval survival, larval size, total adult emergence, and adult emergence timing across the Brassicaceae genotypes (seven *Brassica napus* and a single *Sinapis alba*) selected for larval antibiosis phenotype validation.

| Response | Fixed effect | F | Num. df | Den. df | p |
| --- | --- | --- | --- | --- | --- |
| Larval survival | Genotype | 17.77 | 7 | 59 | **<0.001** |
|  | Plant weight | 6.02 | 1 | 58 | **0.02** |
|  | Block | 3.72 | 2 | 3 | 0.13 |
| Larval size | Genotype | 6.29 | 7 | 59 | **<0.001** |
|  | Block | 2.60 | 2 | 2 | 0.22 |
| Total adults | Genotype | 1.08 | 5 | 52 | 0.38 |
|  | Block | 2.62 | 1 | 52 | 0.11 |
| Weeks to first adult emergence | Genotype | 1.26 | 5 | 52 | 0.29 |
|  | Block | 0.65 | 1 | 52 | 0.42 |
| Mean adult emergence time (weeks) | Genotype | 0.73 | 5 | 52 | 0.61 |
|  | Block | 1.74 | 1 | 52 | 0.19 |

Num. df = numerical degrees of freedom; Den. df = Denominator degrees of freedom (Kenward-Roger estimated for larval survival and larval size responses).

There was also a significant effect of genotype on larval size (mm^2^) (F_(7, 59)_ = 17.77, *p* < 0.001; **Table 2**), with smallest larvae in *S. alba* G1 and largest larvae in *B. napus* 89 (**Figure 3B**). Unlike the diversity screen, larvae were significantly smaller (t_(1, 94)_ = −3.73, *p* < 0.01) in the *S. alba* positive control compared to the negative control (**Figure 3B**), corroborating previous observations of slower larval development in *S. alba*.^56^ Interestingly, we also observed significant differences in larval size between *B. napus* genotypes, with genotypes 84 and 61 hosting smaller larvae than genotype 89 (**84 – 89**: t_(1, 51)_ = −3.90, *p* < 0.01; **61 – 89**: t_(1, 56)_ = −3.81, *p* < 0.01) and no larval size differences between *S. alba* and *B. napus* genotypes 84, 61, and 67 (**Figure 3B**).

All plants in the adult emergence (*n* = 60) experiment showed larval herbivory symptoms. A mean of 6.36±0.30 adult beetles successfully emerged across all plants (equivalent to a 53% larvae-to-adult success rate; **Figure 3C**), with first adult emergence beginning at 8.31±0.08 weeks after infestation (**Figure 3D**). Consistent with the phenotype validation experiment, there was no significant effect of *B. napus* genotype on total adult emergence (F_(5, 52)_ = 1.08, *p* = 0.38), weeks to first adult emergence (F_(5, 52)_ = 1.26, *p* = 0.29; **Table 2**; **Figure 3C-D**), or mean adult emergence time (F_(5, 52)_ = 0.73, *p* = 0.61; **Figure S16**). Per genotype raw means for all variables measured during both experiments can be found in **Tables S9-S13**.

### 3.4. Larval screening can be successfully performed in model species

All 15 infested *B. rapa* R-o-18 plants showed larval herbivory symptoms. Two-weeks post-infestation, a mean of 9.79±0.30 larvae were recovered from the 14 surviving plants (**Figure 4B**). Larvae significantly increased in size (t_(1, 11)_ = 5.19, *p* < 0.001), undergoing a ∼10.5× size increase from 0.45±0.01mm^2^ at hatching to 4.71±0.13mm^2^ as recovered larvae (**Figure 4C**).

**Figure 4.**
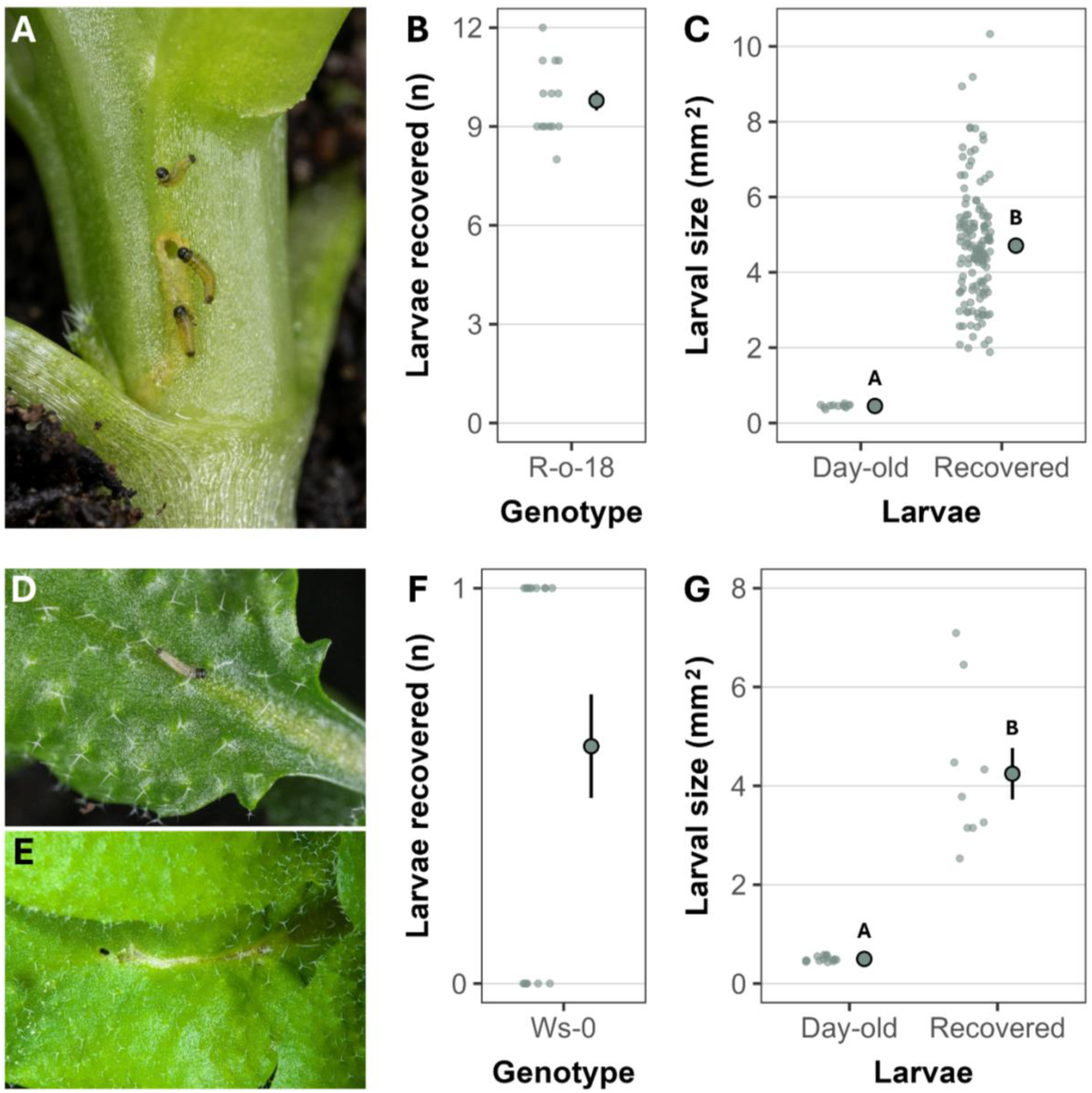
Cabbage stem flea beetle (*Psylliodes chrysocephala*) larval recovery and development within *Brassica rapa* R-o-18 **(A-C)** and *Arabidopsis thaliana* Ws-0 **(D-G)** plants. **(A)** Infestation of *B. rapa* with <24 hour-old larvae. **(B)** Larval recovery from *B. rapa* plants (*n* = 14) two-weeks post-infestation. **(C)** Size (mm^2^) of day-old larvae vs. larvae recovered from *B. rapa* plants. **(D)** Infestation of *A. thaliana* with a single <24 hour-old larva. **(E)** Larval herbivory damage on the midrib of an *A. thaliana* leaf one-week post-infestation. **(F)** Recovery rate (0/1) of larvae from *A. thaliana* plants (*n* = 15). **(G)** Size of <24 hour-old larvae vs. larvae recovered from *A. thaliana* plants. For plots **B**-**C** and **F-G**, large points with whiskers represent the mean ± standard error, left jittered points represent the raw data, and letters represent significant differences (*p* ≤ 0.05) based on larval status.

Fourteen of the 15 infested *A. thaliana* plants (93.33%) presented larval herbivory symptoms and, two-weeks post-infestation, larvae were recovered from nine plants (60%; **Figure 4F**). Larvae significantly increased in size (t_(1, 7)_ = 2.28, *p* = 0.05), developing ∼8.5× in size from 0.50±0.01mm^2^ at hatching to 4.25±0.52mm^2^ as recovered larvae (**Figure 4G**). Successful larval screening within *B. rapa* R-o-18 and *A. thaliana* Ws-0 suggests that the genetic resources available for both models can be exploited to study CSFB larval development and resistance genetics.

## 4. Discussion

CSFB larval herbivory imposes severe yield penalties on OSR growth, with larval antibiosis representing a key breeding target. Despite this, the challenging nature of insect resistance phenotyping has limited attempts to explore CSFB resistance.^56^ Our research aimed to address this by producing a laboratory-based method for controlled larval antibiosis screening,^57^ before: (1) tracking larval development within *B. napus*; (2) exploring larval antibiosis across a large Brassicaceae population; and (3) testing larval screening in two model Brassicaceae relatives – *B. rapa* and *A. thaliana*. We showed that, under our growth conditions, larvae complete development from four-weeks post-infestation, with larval recovery at two-weeks post-infestation providing a faster way of scoring larval antibiosis. Larval screening of the Brassicaceae population revealed limited evidence for resistance within *B. napus* but showed consistent support for larval antibiosis in *S. alba*. Finally, larvae were also able to develop in both *B. rapa* and *A. thaliana*, broadening the genetic resources available to understand CSFB-Brassicaceae interactions.

### 4.1. CSFB larval development

CSFB larvae completed development within *B. napus* from four-weeks post-infestation, as evidenced by a sharp decrease in larval recovery, a 20-fold size increase, and progression through three instars. Given that we did not search for pupae, an alternative explanation may be that larval death occurred at four weeks. However, since later instars are more robust,^43,72^ and adults were first collected at seven-weeks post-infestation during the adult emergence experiment, this seems unlikely. Importantly, larval recovery remained high at two weeks, with larvae significantly increasing in size. Hence, recovery at this timepoint provides a faster method to phenotype larval antibiosis across Brassicaceae populations.

Despite controlled larval infestation, larval size was highly variable within plants. Given variable distribution of glucosinolates across tissues^73,74^, larval development may be influenced by feeding location. Interestingly, we found that larvae avoided apical petioles, where glucosinolates are likely more abundant.^74^ Such patterns have been shown in the generalist *Spodoptera littoralis*^73^ but whether the same effects exist for specialists such as CSFB is unknown. Larval development may also be affected by larval density, especially since CSFB larvae were found to frequently share petioles. Density-dependence is a key driver of insect life-history traits^75,76^ and adverse effects of high density may be stronger in CSFB given that larvae feed internally. Hence, density-dependent effects on CSFB larval development and life-history represent an interesting avenue for future research.

### 4.2. Absence of CSFB larval antibiosis across *Brassica napus*

Our initial larval diversity screen identified weak genotype effects on larval survival across the *B. napus* population. However, in follow-up experiments with larger per-genotype sample sizes, differences between initially identified ‘resistant’ and ‘susceptible’ genotypes were no longer present, with similar larval survival, larval size, adult emergence, and emergence timing found across genotypes. Importantly, the screened *B. napus* population shows variability across other traits.^58–60^ Hence, if CSFB larval antibiosis did exist within this panel we should have been able to identify it. This is a crucial finding for the field, since researchers and breeders alike invest substantial resources trying to identify larval antibiosis under field conditions. However, if we cannot find larval antibiosis across a large population with highly controlled phenotyping approaches, it seems unlikely that field trials will provide any promising candidates. A lack of CSFB larval antibiosis in *B. napus* was also shown in Döring & Ulber,^56^ albeit within a much smaller population (four genotypes versus 97 in the present study). Further, *B. napus* appears to lack resistance to multiple insect pests,^23,38^ possibly due to limited genetic diversity after hybridisation followed by selection against glucosinolates during oilseed domestication.^25,32–35^ All genotypes examined in this study have undergone domestication but, given the inclusion of swedes and kales, not necessarily glucosinolate-based selection. Hence, it is possible that insect resistance traits besides glucosinolate production have been lost during *B. napus* domestication. However, a thorough comparative investigation between *B. napus* and insect-resistant relatives or feral *Brassica* species is required to confirm this.

Our study prioritised larval survival over larval size as a measure of larval antibiosis. This was because our phenotyping method involved larval recovery using the Berlese method.^61^ As a result, we were able to screen large numbers of plants at the expense of accurate larval size, given larvae can continue developing within plants before recovery.^61^ We are unable to distinguish between larval death within the plant versus larvae failing to penetrate the plant. However, given that all infested plants showed signs of larval herbivory symptoms on the final day, and desiccated larvae were never found on the plant exterior, we are confident that larval survival represents mortality experienced within the plant. Larval antibiosis may take place later than two-weeks post-infestation but, given that antibiosis within *S. alba* was consistently observed at two weeks, we deem this unlikely.

In the present study, we only assessed the presence of larval antibiosis across *B. napus* genotypes. However, alternative measures of CSFB resistance such as antixenosis and tolerance^18,21,22^ to both adults and larvae across *B. napus* diversity remain untested. For instance, oviposition preference differences across Brassicaceae species have been shown in multiple insects, including cabbage butterflies (*Pieris brassicae* and *P. rapae*),^77^ cabbage seedpod weevil *(Ceutorhynchus obstrictus),*^78^ and pollen beetle (*Meligethes aeneus*).^79^ Oviposition preferences for CSFB are currently unknown; however, oviposition rate may depend on the quality of the host plant and availability of digestible carbohydrates.^80^ Further, tolerance to low larval loads (infestation of less than five larvae per plant) has been demonstrated in a single *B. napus* genotype, with yield declines observed with increasing larval density.^43^ However, per-genotype relationships between larval density and plant yield are unexplored. Hence, while larval antibiosis is unlikely to exist with *B. napus*, alternative mechanisms of plant resistance may yet be discovered.

### 4.3. Brassicaceae relatives can inform CSFB larval resistance

*Sinapis alba*, a relative of *B. napus*, consistently demonstrated CSFB larval antibiosis, with lower larval survival and smaller larvae compared to the screened *B. napus* genotypes. Promisingly, CSFB larval antibiosis has also been reported in a separate *S. alba* genotype,^56^ suggesting that larval resistance may be a shared trait across *S. alba* genetic diversity. Future work should look to confirm whether larval antibiosis is the rule within *S. alba* by applying our screening methodology^57^ to a genetically diverse population. Additionally, since we only measured larval survival and size at two-weeks post-infestation, it would be interesting to see if adults can successfully emerge from *S. alba* and whether adults suffer any adverse effects, such as reduced fecundity or lifespan.^81^ *Sinapis alba* shows resistance to a range of other insect pests,^82–84^ and, in some cases, this resistance has been successfully tracked to QTLs and introgressed into OSR.^85,86^ For instance, a mapping population derived from a resistant × susceptible *S. alba* cross allowed the identification of a QTL explaining 10% of heritable variation associated with cabbage seedpod weevil (*C. obstrictus*) resistance.^85^ Hence, identifying the metabolic and genetic factors associated with CSFB resistance in *S. alba* and introgressing these traits into *B. napus* may represent a key strategy in breeding CSFB-resistant OSR.

In the present study, we showed that CSFB larvae can successfully infest and develop within both *B. rapa* R-o-18 and *A. thaliana* Ws-0. Alongside *S. alba*, both *B. rapa* and *A. thaliana* may also provide insights into CSFB resistance. As the progenitors of *B. napus*, genetic diversity is far higher in both *B. rapa* and *B. oleracea*,^25^ and both species may contain sources of CSFB resistance. An interesting approach could be to screen populations of both species for CSFB resistance before performing crosses to pyramid genes and produce resynthesised, resistant *B. napus* genotypes. Such approaches have previously been applied to resynthesise *B. napus* with resistance to the fungal pathogen *Verticillium longisporum*,^87,88^ while resynthesised *B. napus* has also shown increased resistance to rape stem weevil (*C. napi*).^89^ In addition, *B. rapa* R-o-18 offers a selection of TILLING mutants,^64^ providing a rich resource for testing gene function in relation to larval resistance.

CSFB larval screening in *Arabidopsis* also offers a promising alternative to identify resistance genes. Alongside the huge genetic diversity available in *Arabidopsis*, larval screening could be far higher throughput than in *Brassica* species, given the requirement for fewer larvae (one larva per plant in *Arabidopsis* versus 12 in *Brassica*), fewer infestation days (one day in *Arabidopsis* versus four in *Brassica*), and less space. Plant-insect association studies across diverse *Arabidopsis* populations have identified candidate genes for cabbage butterfly (*P. brassicae*) oviposition,^90^ cabbage whitefly (*Aleyrodes proletella*) survival and fecundity,^91^ and diamondback moth (*Plutella xylostella*) larval herbivory.^92^ Such approaches could therefore be used to identify CSFB resistance candidates in *Arabidopsis* before searching for homologs within *Brassica* genomes. Further, the rich diversity of knock-out and over-expression lines in *Arabidopsis* offers a convenient method for testing CSFB resistance gene function.

### 4.4 Conclusions

We outline a method for highly controlled CSFB larval screening within Brassicaceae and, by applying this method to a diverse collection of Brassicaceae genotypes, show limited evidence for larval antibiosis within *B. napus* but consistent antibiosis in *S. alba*. These findings highlight the need for researchers and breeders to look beyond *B. napus* genetic diversity to uncover genes involved in CSFB resistance, with Brassicaceae relatives representing a promising resource. Additionally, the suitability of *B. rapa* and *A. thaliana* for larval screening provides further genetic tools to mine for CSFB resistance. Finally, beyond the use of these methods for gene and trait discovery in OSR breeding, we also suggest that the CSFB-Brassicaceae system could provide fundamental insights into plant-insect interactions for insects with larval mining strategies.

## 5. Acknowledgements

We thank Fred Stevenson (JIC) for assistance with larval imaging; Mark Nightingale (Elsoms) for providing *S. alba* G1; Dr Samantha Cook and Dr Patricia Ortega-Ramos (Rothamsted Research) for feedback on methodology and results; Dr Emmanuel Solomon (JIC) for *A. thaliana* growth; Phil Robinson (JIC) for larval infestation photography; the JIC Entomology platform for support throughout; and all members of the Wells group for feedback on an earlier version of this manuscript.

This work was supported by a UKRI BBSRC grant awarded to RW and SP (BB/V015524/1) and a Royal Society of Biology Plant Health Undergraduate Studentship grant awarded to REB. RW and SP acknowledge support from BBSRC Institute Strategic Programme funding (APH – BB/X010996/1 and BRiC – BB/X01102X/1).

## 6. Author contributions

**Ryan E. Brock:** conceptualisation (supporting); methodology (lead); investigation (lead); data curation (lead); formal analysis (lead); visualisation (lead); funding acquisition (supporting); supervision (supporting); writing – original draft (lead); writing – review and editing (equal).

**Clara Courtney:** methodology (supporting); investigation (supporting); writing – review and editing (supporting).

**Steven Penfield:** conceptualisation (supporting); funding acquisition (supporting); supervision (supporting); writing – review and editing (supporting).

**Rachel Wells:** conceptualisation (lead); funding acquisition (lead); supervision (lead); writing – review and editing (equal).

## 8. Data availability statement

All data and code required for visualisation and analyses are available from Zenodo (DOI: 10.5281/zenodo.18009384).^93^

