## Supplementary materials for "Larval antibiosis to cabbage stem flea beetle (*Psylliodes chrysocephala*) is absent within oilseed rape (*Brassica napus*)"

**Supplementary methods**

**1. Model fitting for larval development time course analyses**

We examined cabbage stem flea beetle (*Psylliodes chrysocephala*; CSFB) larval development within *Brassica napus* over time by fitting total live larvae recovered, larval size, and larval instar proportion as a function of timepoint (**Models 1-3** in **Table S3**). The larval recovery model also contained an effect for plant weight at dissection (since larger plants may support more larvae), while the larval size model contained a random effect for nesting of larvae within samples. Larval instar proportions were fitted as a response by first producing a mean instar score per plant, calculated as:

*(total L1 larvae) + (2 × total L2 larvae) + (3 × total L3 larvae)*

*total larvae*

We assessed changes in larval size across instars by fitting larval size as a function of instar, with a random effect for nesting of larvae within plants (**Model 4** in **Table S3**).

The larval size model included the 20 <24 hour-old larvae collected during infestation (i.e. the 0W timepoint), while the larval size by instar model included only those larvae recovered from dissected plants. Sample sizes for all models can be found in **Table S3**.

**2. Model fitting for Brassicaceae population larval screening analyses**

We defined CSFB larval antibiosis across Brassicaceae genotypes as both the number of larvae recovered (i.e. larval survival) and the size of recovered larvae (i.e. larval development). We examined differences in larval survival and development across genotypes by fitting larval recovery and larval size as a function of genotype (**Models 5-6** in **Table S3**).

The maximal larval recovery model also included plant weight, replicate, and shelf as fixed effects, and block (nested within replicate) and shelf position (nested within shelf) as random effects. Both shelf and shelf position were dropped from the larval recovery model, due to non-significance and singular fit, respectively. The maximal larval size model also included plant weight (since larger plants may support larger larvae), total larvae recovered from the plant (to account for possible density-dependent effects on larval size), replicate, and shelf as fixed effects, and block, shelf position, and plant ID (for nesting of larvae within plants) as random effects. Plant weight, total larvae recovered, and shelf effects were all dropped from the larval size model due to non-significance. Despite non-significance in both models, the replicate effect was retained to reflect experimental design.

We also explored per genotype mean larval size as a function of mean larval recovery (i.e. whether genotypes with higher larval recovery also have larger larvae), using both raw means and estimated means fitted from the larval recovery and larval size models (**Models 7-8** in **Table S3**).

**3. Model fitting for phenotype validation and adult emergence analyses**

For phenotype validation, larval antibiosis across genotypes was defined as before, with two extra measures of antibiosis in the adult emergence experiment: total adult emergence (i.e. the number of larvae that successfully completed development) and time to first adult emergence (i.e. larval development time).

We explored differences in larval survival and development by fitting larval recovery and larval size as a function of plant genotype (**Models 9-10** in **Table S3**). The maximal larval recovery model also included plant weight, block, and shelf as fixed effects, and shelf position (nested within shelf) as a random effect. The shelf fixed effect was dropped from the larval recovery model due to non-significance. The maximal larval size model also included plant weight, total larvae recovered from the plant, block, and shelf as fixed effects, and plant ID (for nesting of larvae within plants) and shelf position as random effects. Fixed effects of plant weight, total larvae recovered, and shelf were all dropped from the larval size model due to non-significance. Despite non-significance in both models, the block effect was retained to reflect experimental design.

We assessed differences in adult emergence and emergence time by fitting total adults, weeks to first adult emergence, and mean adult emergence time as a function of genotype (**Models 11-13** in **Table S3**). Weeks to first adult emergence was calculated for each plant as:

*date of first adult emergence – date of first larval infestation day*

*7*

Mean adult emergence time was calculated for each plant as:

*weeks to first adult emergence + (date of last adult emergence – date of first adult emergence)*

*2*

All models also included block as a fixed effect and shelf position as a random effect. Shelf was not included as a fixed effect within these models, since shelf was synonymous with block (i.e. all plants within a block were all grown on the same shelf **Figure S6**). Due to singular fit, the shelf position random effect was dropped from all models, leading to linear models for the three responses rather than linear mixed models. In all models, block was retained as an effect despite non-significance to reflect experimental design.

**4. Model fitting for Brassicaceae relative larval screening analyses**

For both *Brassica rapa* R-o-18 and *Arabidopsis thaliana* Ws-0, we explored larval development within plants by fitting larval size as a function of status, where status had two levels: day-old (the size of larvae when introduced to the plant) and recovered (the size of larvae when recovered from the plant through evacuation). Both models also included a random effect for nesting of larvae within samples (**Models 14-15** in **Table S3**).

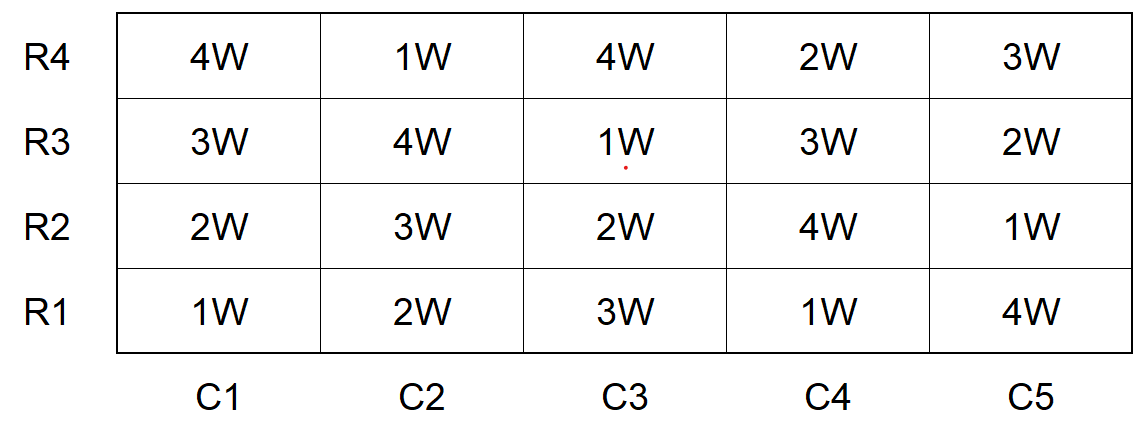
**Figure S1.** Layout for the cabbage stem flea beetle (*Psylliodes chrysocephala*; CSFB) larval development time course experiment performed in a single *Brassica napus* genotype (Apex-93_5 × Ginyou_3 DH). Twenty plants infested with 12 CSFB larvae each were grown in a 4×5 row-column matrix, with each column containing a single replicate of each dissection timepoint: 1W = plant dissected one week after larval infestation; 2W = plant dissected two weeks after larval infestation; 3W = plant dissected three weeks after larval infestation; 4W = plant dissected four weeks after larval infestation.

**Supplementary figures**

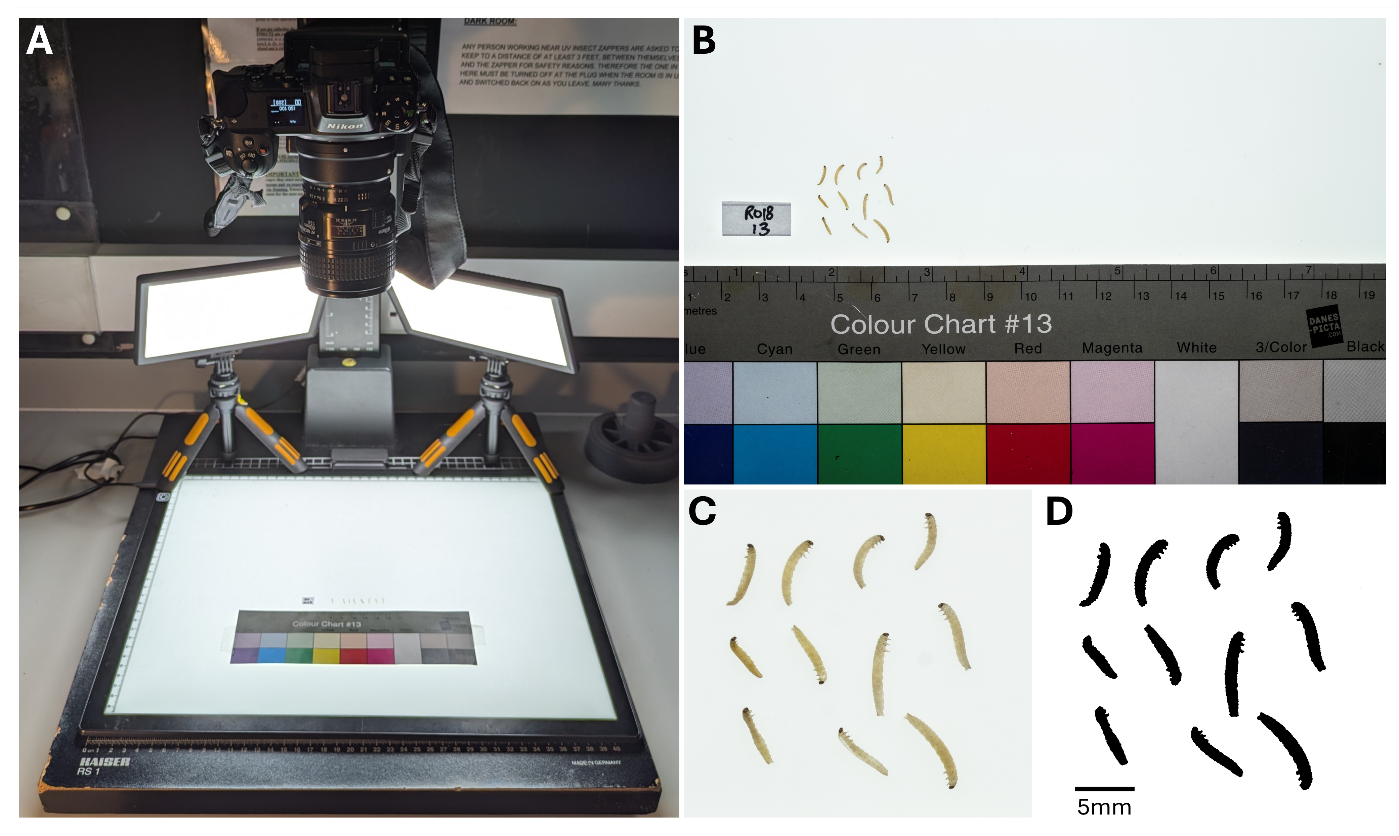
**Figure S2.** Setup for imaging and measuring cabbage stem flea beetle (*Psylliodes chrysocephala*; CSFB) larvae recovered from Brassicaceae plants (*Brassica napus*, *B. rapa*, *Sinapis alba*, and *Arabidopsis thaliana*). **(A)** Photography setup comprising an overhead camera mount (Kaiser RS1, Kaiser Fototechnik, Buchen, Germany), DSLR camera (Nikon Z6 with AF Micro Nikkor 60mm lens, Nikon, Tokyo, Japan), A3 LED light pad (AGM, Manchester, UK), two overhead LED lights (Raleno PLV-S116, KEJI, Shenzen City, China), and a scale/colour chart (Wex Photo Video, Norwich, UK). **(B)** Raw image of CSFB larvae recovered from an infested *B. rapa* R-o-18 plant, including label (genotype + replicate number) and scale. **(C)** Cropped raw image, containing only the larvae to be measured. **(D)** Binary conversion of larval image, allowing measurement of larval size (in mm^2^) using the ‘Analyse Particles’ function in ImageJ v1.52a. Scale bar represents 5mm for **C** and **D**.

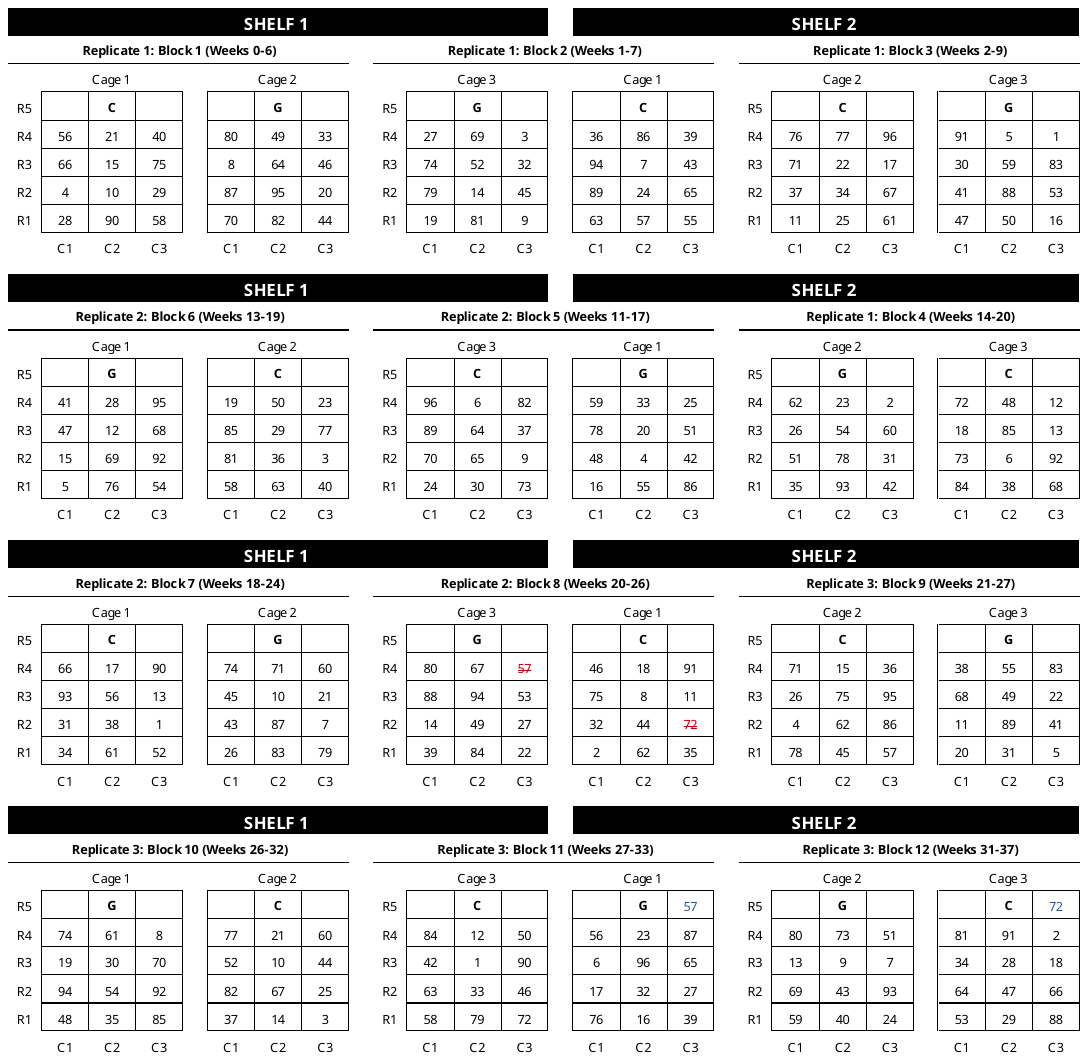
 **Figure S3.** Block layouts for phenotyping 96 *Brassica napus* genotypes for cabbage stem flea beetle (*Psylliodes chrysocephala*) larval antibiosis. *B. napus* genotypes were randomised across a temporal 12-block alpha design (plots per block (*k*) = 24, replicates per genotype (*r*) = 3, blocks per replicate (*s*) = 4), and grown across two CER shelves, with a new block beginning every 1-9 weeks. Every block also contained a negative *B. napus* control (KWS Campus = C) and a positive *Sinapis alba* control (Elsoms G1 = G). Each block was split across two cages, with each cage setup in a 5×3 row-column matrix. For block 8, two genotypes (57 and 72) died prior to larval infestation (cells with red strikethrough text). These genotypes were therefore placed into later blocks (57 into block 11 and 72 into block 12; cells with blue text). Each number (1-96) represents a *B. napus* genotype from the RIPR diversity set (**Table S1**).

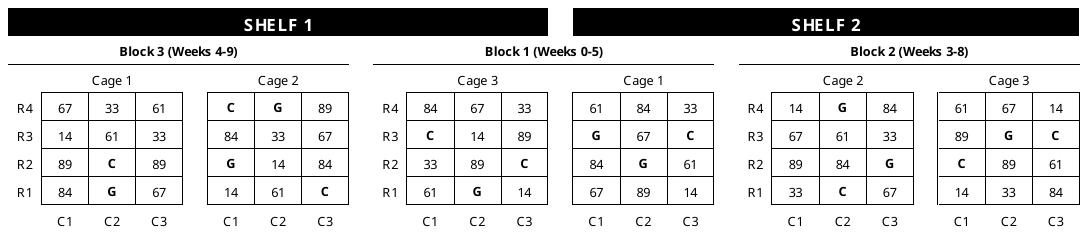
**Figure S4.** Block layouts for screening eight Brassicaceae genotypes (seven *Brassica napus* and a single *Sinapis alba*; **Table S1**) for cabbage stem flea beetle (*Psylliodes chrysocephala*) larval antibiosis phenotype validation. Genotypes included the ‘resistant’ *B. napus* (genotypes 33, 61 and 67), the ‘susceptible’ *B. napus* (genotypes 14, 84 and 89), the negative *B. napus* control (KWS Campus = C), and the positive *S. alba* control (Elsoms G1 = G). Genotypes were randomised across three blocks, with a new block starting every 1-3 weeks. All blocks were grown across two CER shelves, with each block containing three replicates of each genotype. Each block was split across two cages, with each cage set up in a 4×3 row-column matrix.

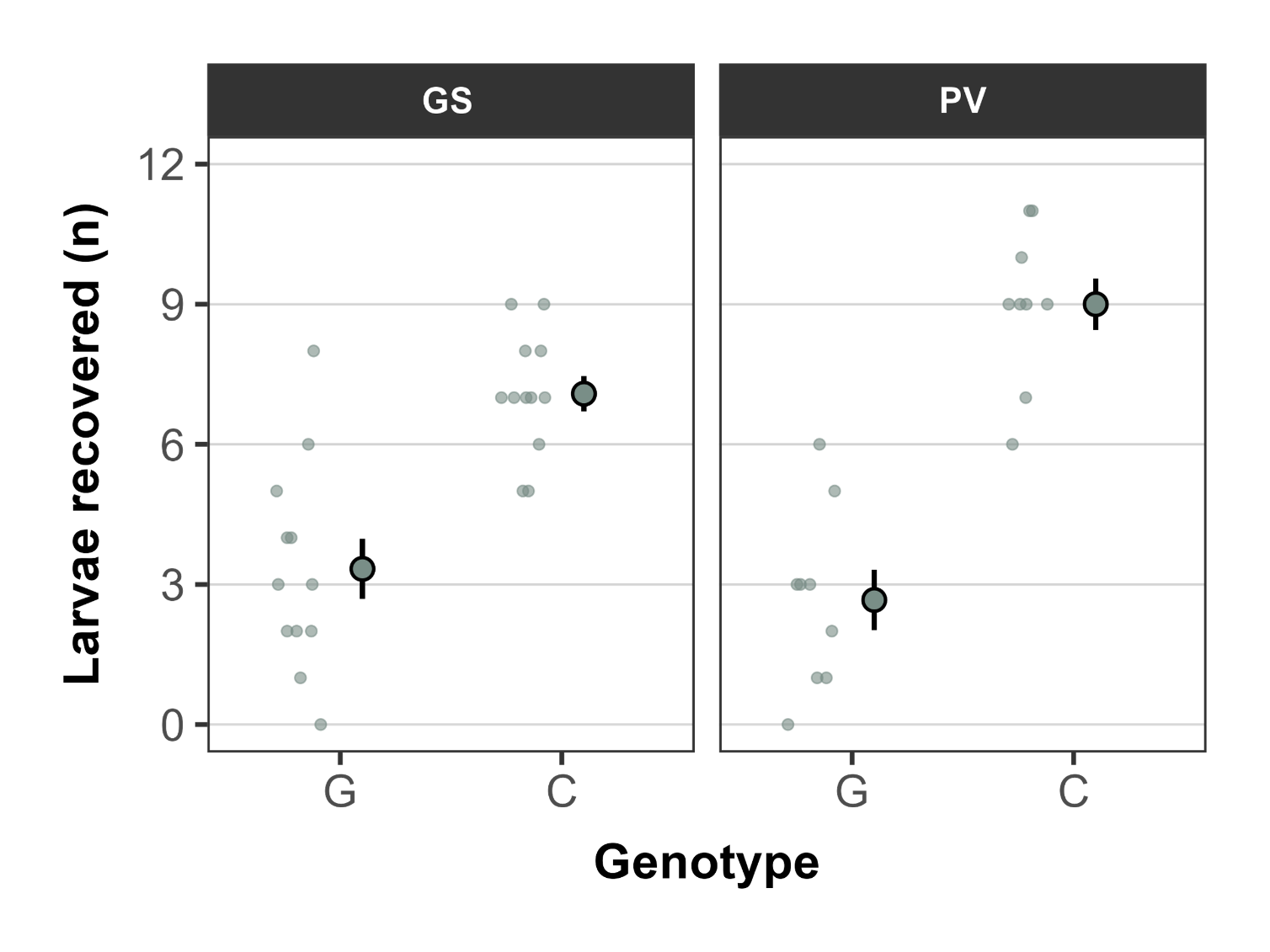
**Figure S5.** Comparison of cabbage stem flea beetle (*Psylliodes chrysocephala*) larval survival in the two control genotypes (G = *Sinapis alba* Elsoms G1; C = *Brassica napus* KWS Campus) between the Brassicaceae genotype screening experiment (GS, left) and phenotype validation experiment (PV, right). Larval survival is defined as the number of CSFB larvae recovered from a plant two weeks after infestation with 12 larvae. Large points with whiskers represent the mean ± standard error and jittered points to the left represent the raw data. Statistical analyses were not run to compare differences in larval survival between the two experiments given that the independent experiments were run with different designs (**Figure S3-S4**).

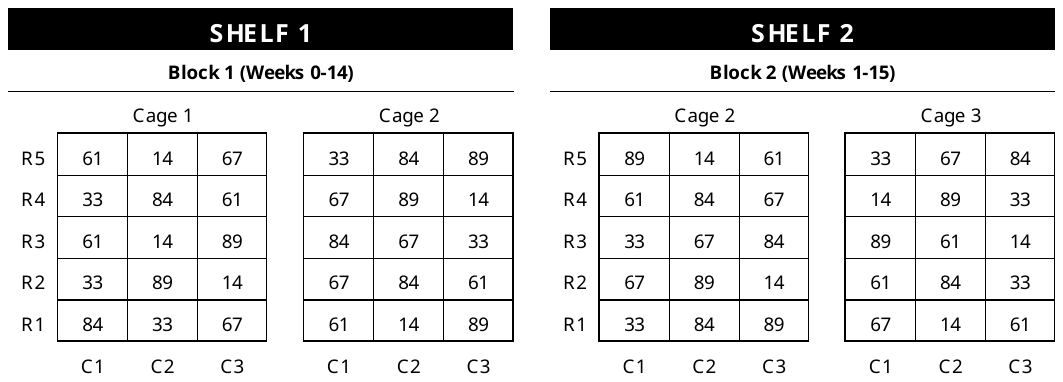
**Figure S6.** Block layouts for screening six *Brassica napus* genotypes for cabbage stem flea beetle (*Psylliodes chrysocephala*) larval antibiosis (via adult beetle emergence). Genotypes included the ‘resistant’ *B. napus* (genotypes 33, 61 and 67), and the susceptible *B. napus* (genotypes 14, 84 and 89). Genotypes were randomised across two blocks, with a new block starting each week. Both blocks were grown across two CER shelves, with each block containing five replicates of each genotype. Each block was split across two cages, with each cage set up in a 5×3 row-column matrix. Full genotype details can be found in **Table S1**.

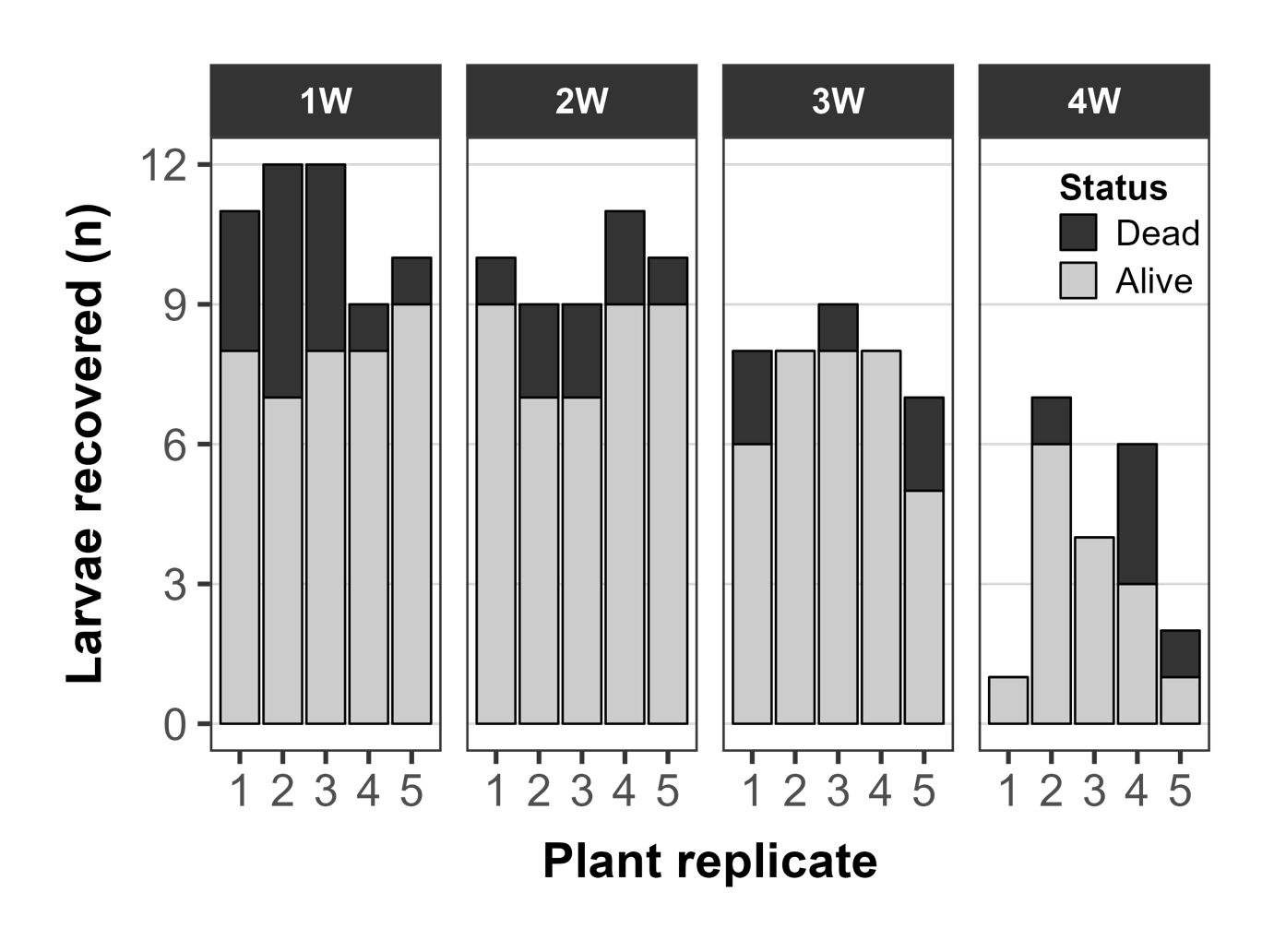
**Figure S7.** Total recovery of cabbage stem flea beetle (*Psylliodes chrysocephala*) larvae from *Brassica napus* (Apex-93_5×Ginyou_3 DH) plants infested with 12 larvae each. Plants were dissected over four timepoints (1-4 weeks) following infestation, with five plants dissected per timepoint. Grey bars = total live larvae found within each plant; black bars = total dead larvae found within each plant.

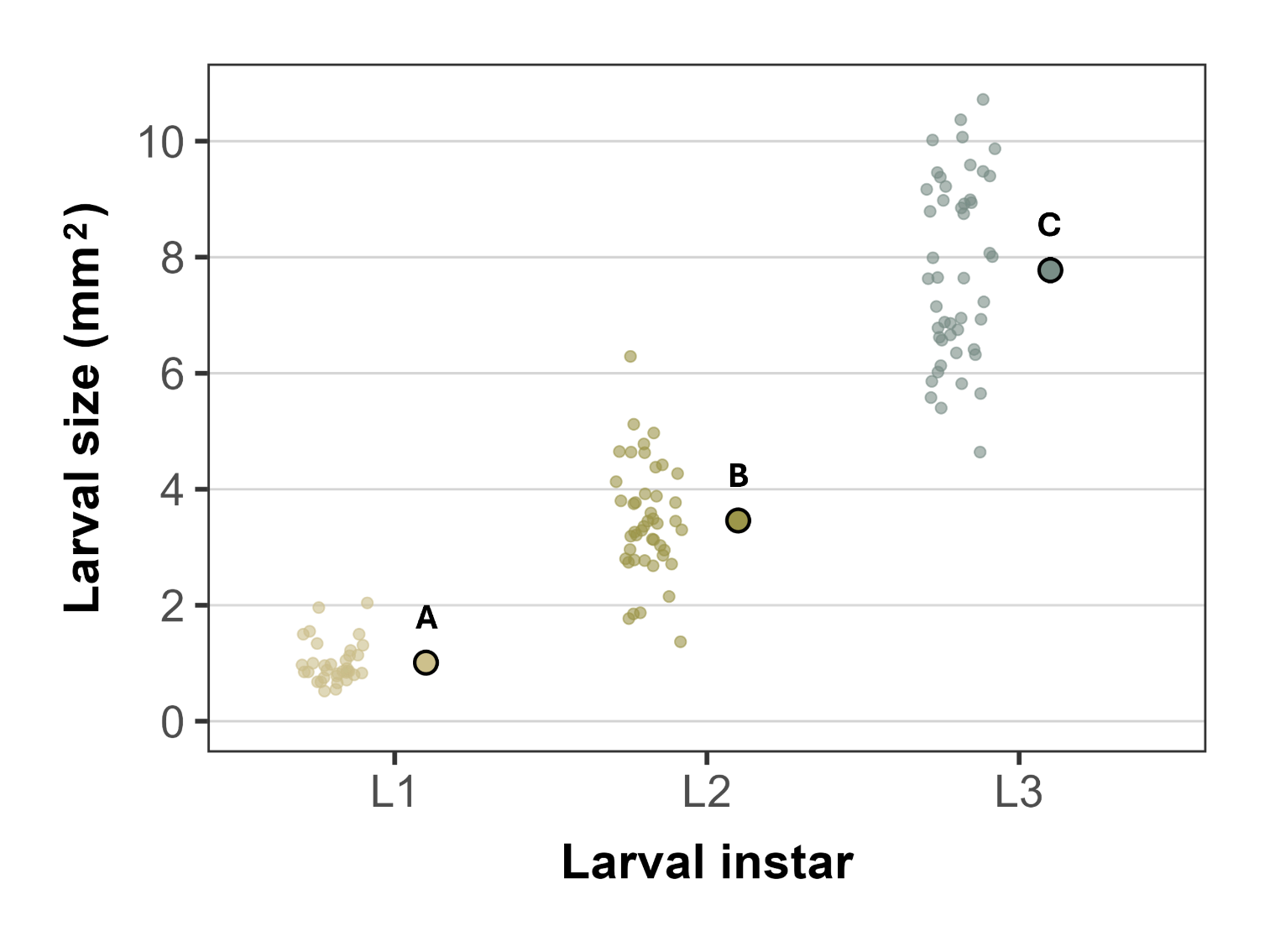
**Figure S8.** Size of cabbage stem flea beetle (*Psylliodes chrysocephala*; CSFB) larvae recovered from infested *Brassica napus* (Apex-93_5×Ginyou_3 DH) plants by larval instar. Large points with whiskers represent the mean ± standard error, jittered points to the left represent the raw data, and letters represent significant differences (*p* < 0.05) between larval instars.

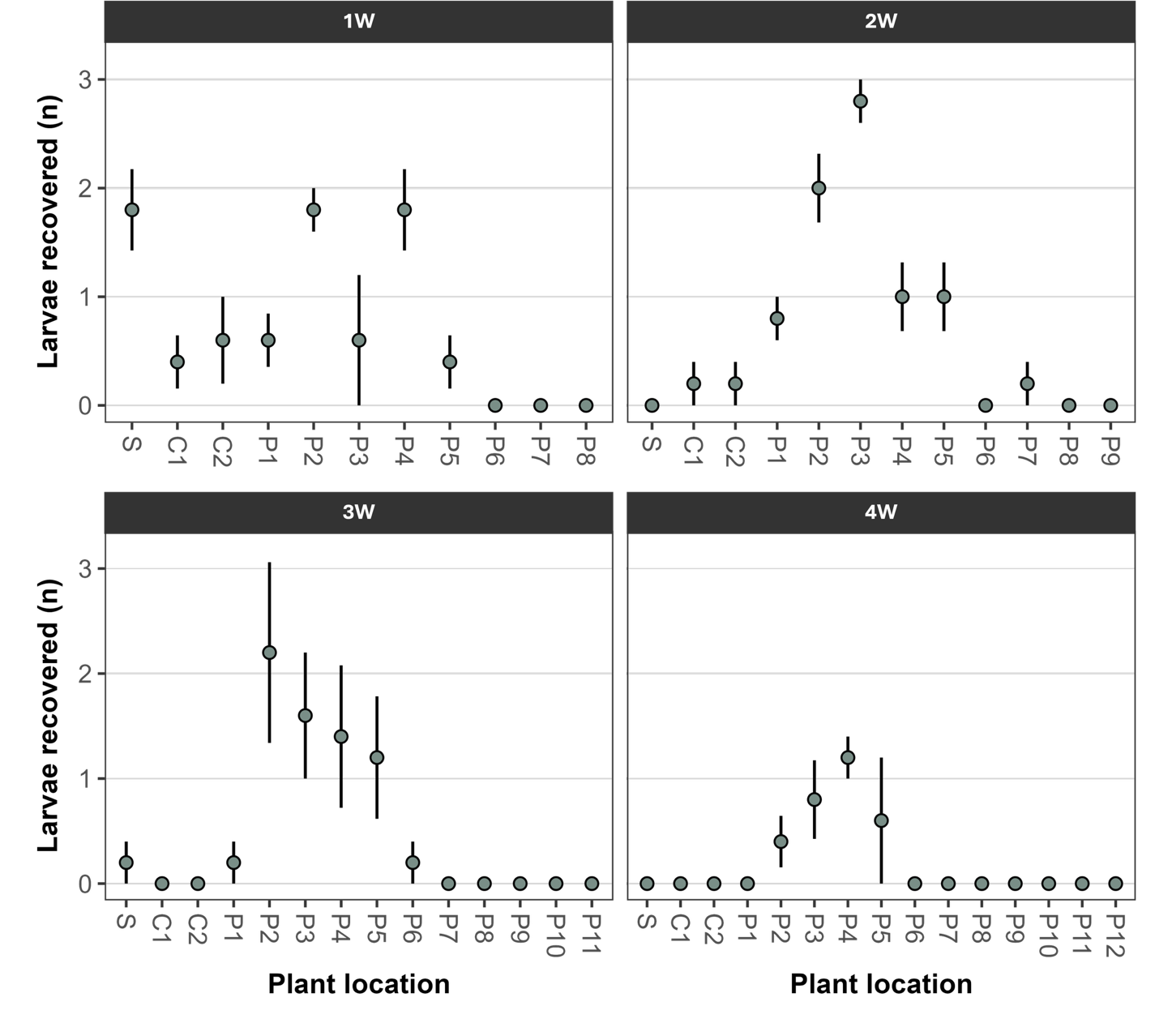
**Figure S9.** Location of cabbage stem flea beetle (*Psylliodes chrysocephala*) larvae recovered from *Brassica napus* (Apex-93_5×Ginyou_3 DH) plants infested with 12 larvae each. Plants were dissected over four timepoints (1-4 weeks) following infestation, with five plants dissected per timepoint. S = stem; C1-2 = cotyledons; P1-12 = true leaf petioles, with P1 representing the first true leaf petiole and sequential numbers representing the next true leaf (i.e. high P numbers represent the youngest true leaf petioles, closest to the plant apex). Large points with whiskers represent the mean ± standard error.

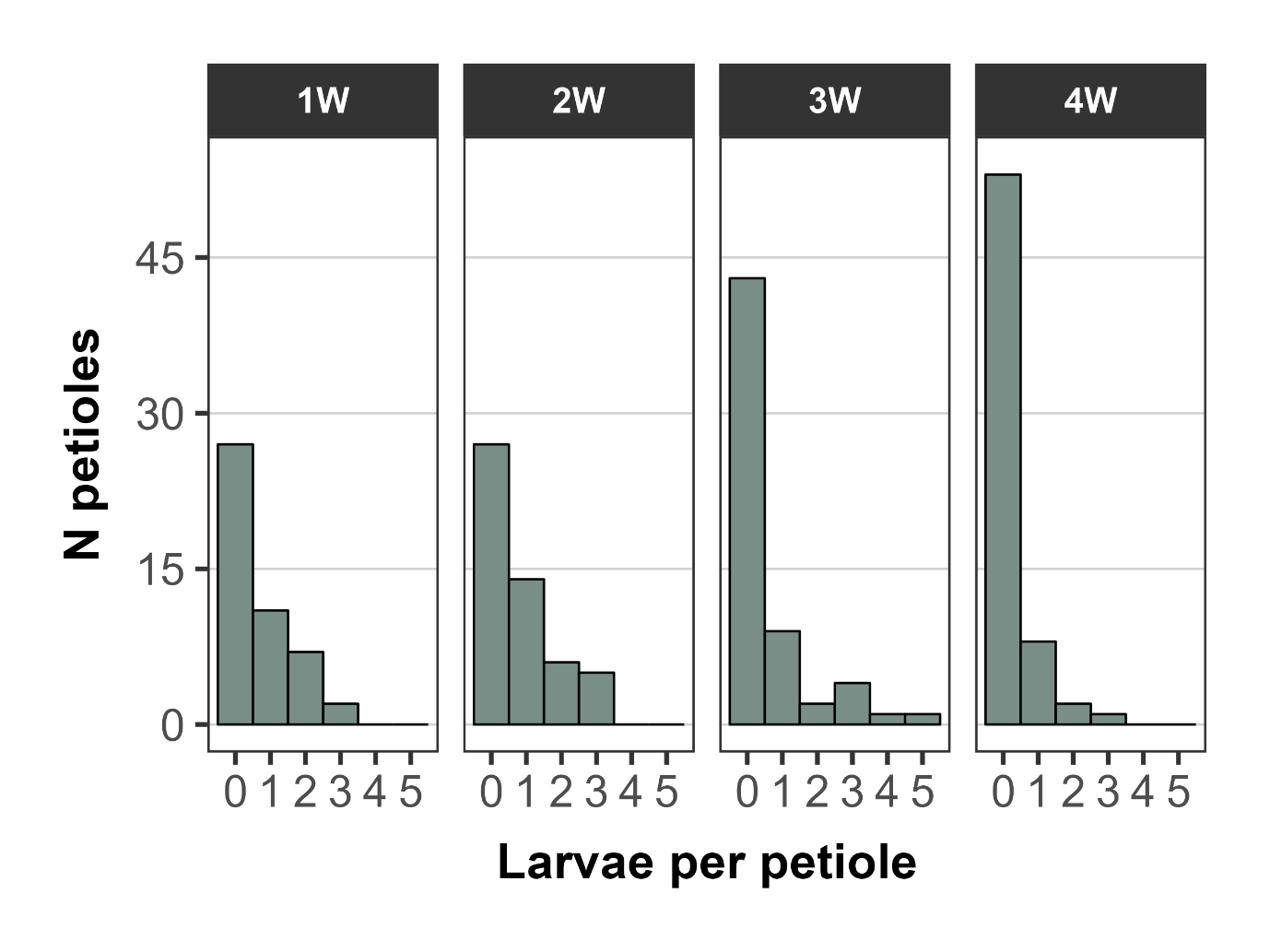
**Figure S10.** Number of cabbage stem flea beetle (*Psylliodes chrysocephala*) larvae per petiole in *Brassica napus* (Apex-93_5×Ginyou_3 DH) plants infested with 12 larvae each. Plants were dissected over four timepoints (1-4 weeks) following infestation, with five plants dissected per timepoint. Hence, bars within each timepoint represent the total number of dissected petioles across all plants within a timepoint. Occurrences of two or more larvae per petiole are defined as petiole sharing.

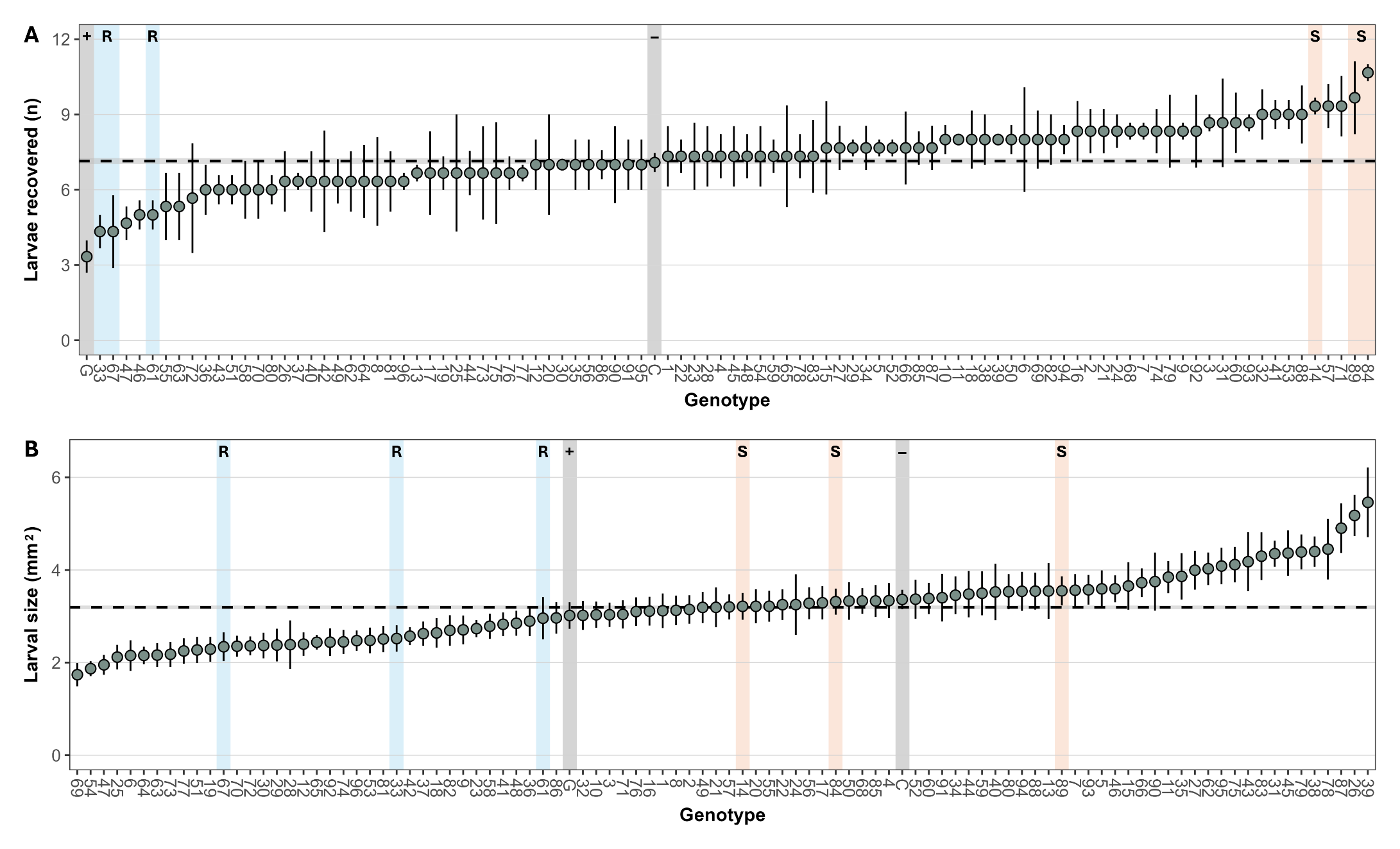
 **Figure S11.** Per genotype mean ± standard error for cabbage stem flea beetle (*Psylliodes chrysocephala*) larval survival **(A)** and larval size **(B)** across 98 Brassicaceae genotypes (97 *Brassica napus* and a single *Sinapis alba*; **Table S1**). Larval survival is defined as the number of CSFB larvae recovered from a plant two weeks after infestation with 12 larvae. For both plots, genotypes are arranged from lowest to highest mean. The dashed horizontal line with grey shading represents the mean ± standard error for all data. Blue and peach highlighted bars represent genotypes identified as resistant (R) and susceptible (S), respectively. Grey highlighted bars represent the positive (G = *S. alba* Elsoms G1) and negative (C = *B. napus* KWS Campus) controls, respectively.

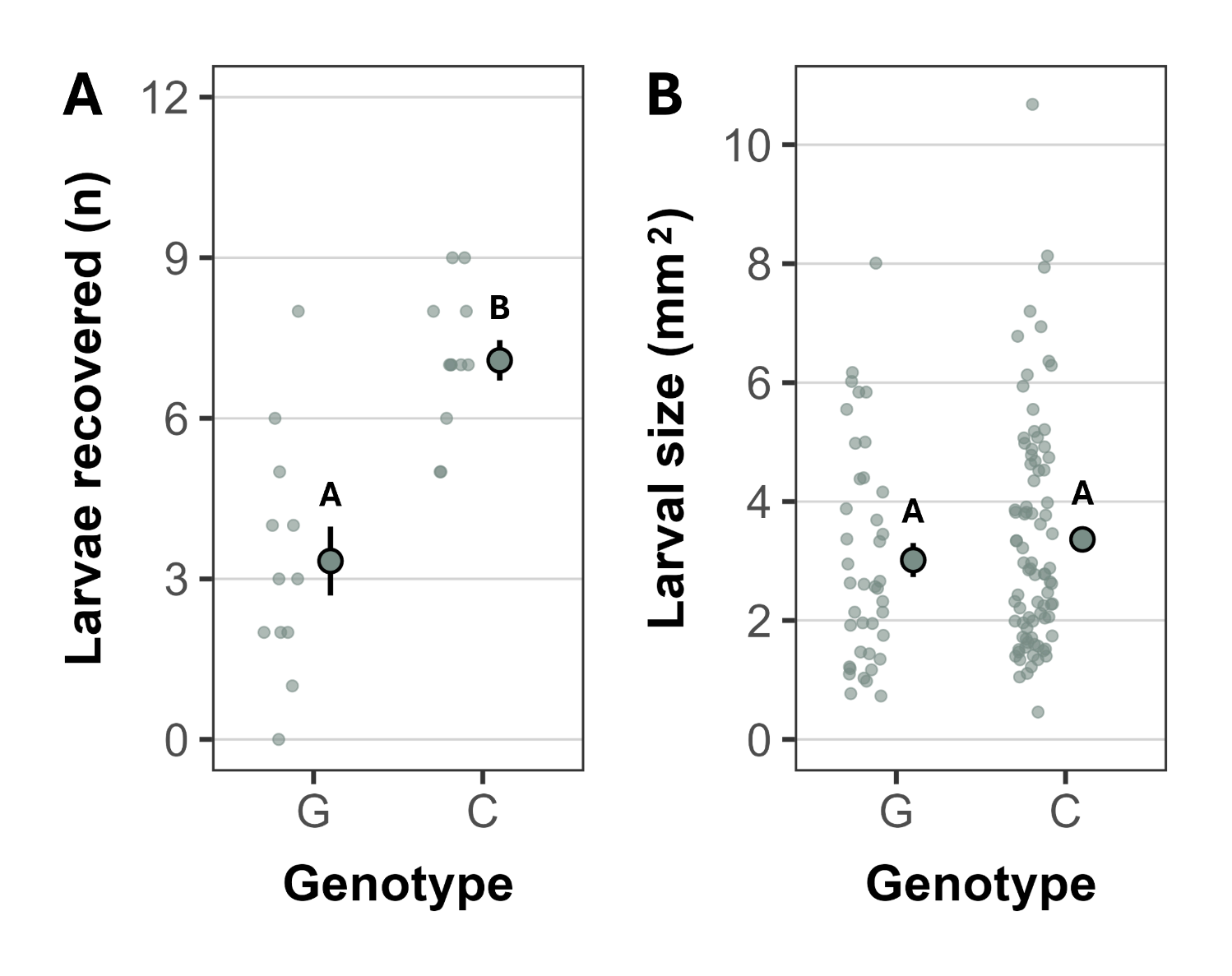
**Figure S12.** Cabbage stem flea beetle (*Psylliodes chrysocephala*) larval survival **(A)** and larval size **(B)** in the two control genotypes (G = *Sinapis alba* Elsoms G1; C = *Brassica napus* KWS Campus) used for the *B. napus* population larval antibiosis screen. Larval survival is defined as the number of CSFB larvae recovered from a plant two weeks after infestation with 12 larvae. Large points with whiskers represent the mean ± standard error, jittered points to the left represent the raw data, and letters represent significant differences (*p* < 0.05) between genotypes.

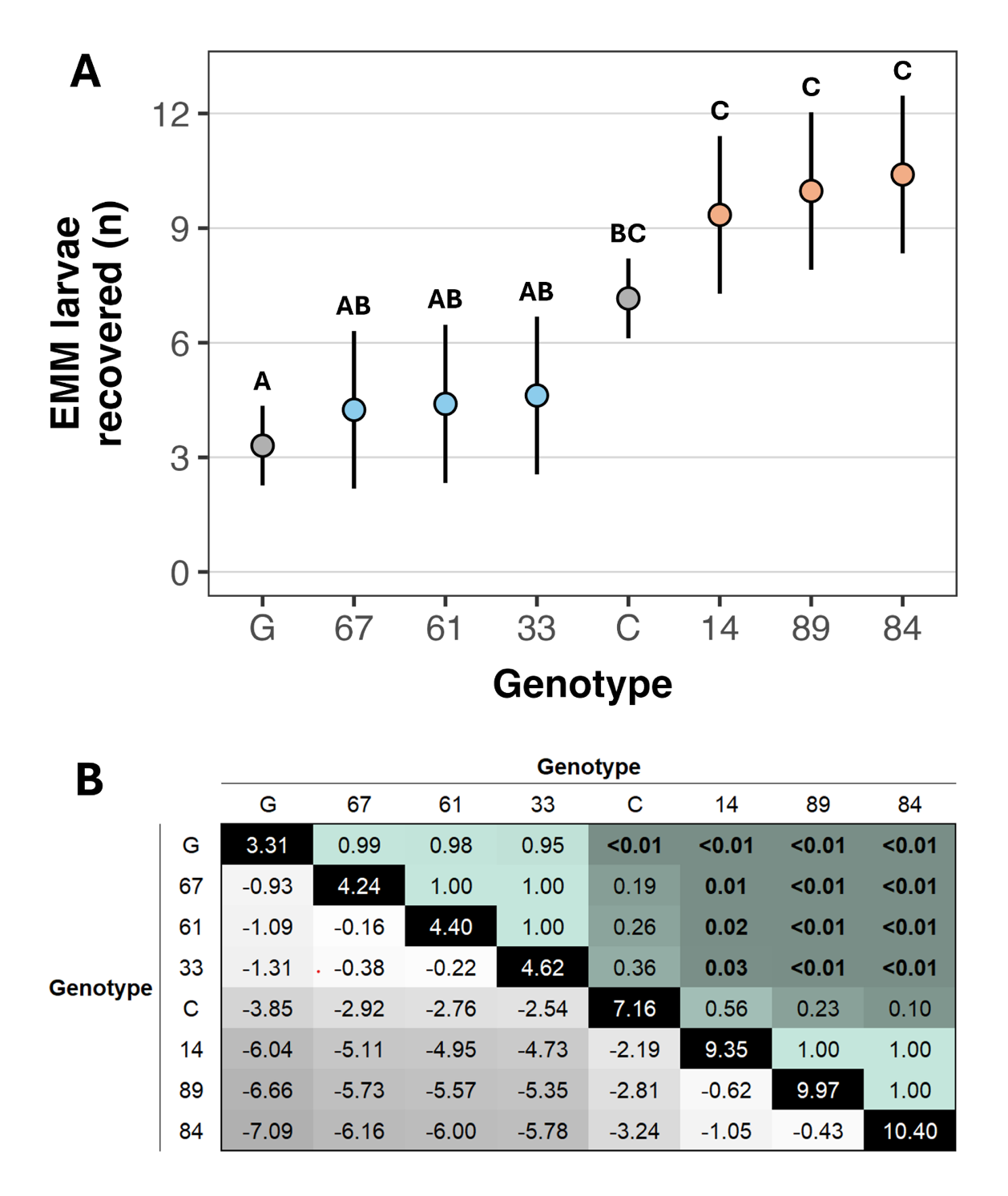
 **Figure S13.** Per genotype estimated marginal means and contrasts for cabbage stem flea beetle (*Psylliodes chrysocephala*; CSFB) larval survival across eight selected Brassicaceae genotypes (seven *Brassica napus* and a single *Sinapis alba*; **Table S1**) from the Brassicaceae population larval antibiosis screen. Larval survival is defined as the number of CSFB larvae recovered from a plant two weeks after infestation with 12 larvae. **(A)** Per genotype EMM ± estimated 95% confidence intervals for larval survival across resistant (blue), susceptible (peach), and control (grey) plant genotypes (G = *S. alba* Elsoms G1; C = *B. napus* KWS Campus). Letters represent significant differences (*p* < 0.05) between genotypes. **(B)** Contrast matrix for larval survival across the eight genotypes. Values in the black diagonal boxes represent EMMs for each genotype, values in the left-hand triangle represent EMM contrasts between genotypes (darker grey = larger contrast), and values in the right-hand triangle represent significance thresholds between genotype comparisons (darker green = smaller *p* value).

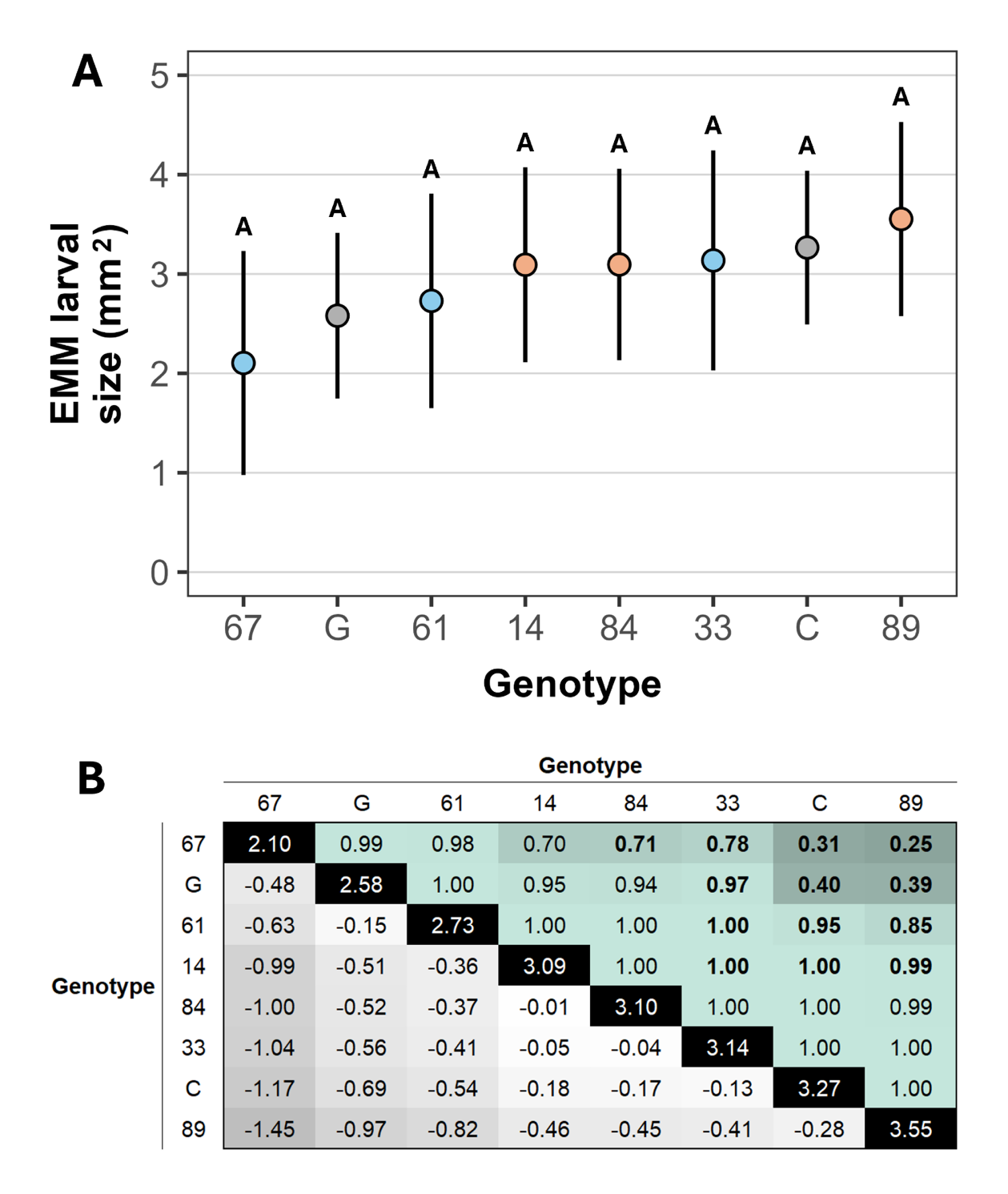
**Figure S14.** Per genotype estimated marginal means and contrasts for cabbage stem flea beetle (*Psylliodes chrysocephala*; CSFB) larval size across eight selected Brassicaceae genotypes (seven *Brassica napus* and a single *Sinapis alba*; **Table S1**) from the Brassicaceae population larval antibiosis screen. **(A)** Per genotype EMM ± estimated 95% confidence intervals for larval size across resistant (blue), susceptible (peach), and control (grey) plant genotypes (G = *S. alba* Elsoms G1; C = *B. napus* KWS Campus). Letters represent non-significant differences (*p* > 0.05) between genotypes. **(B)** Contrast matrix for larval size across the eight genotypes. Values in the black diagonal boxes represent EMMs for each genotype, values in the left-hand triangle represent EMM contrasts between genotypes (darker grey = larger contrast), and values in the right-hand triangle represent significance thresholds between genotype comparisons (darker green = smaller *p* value).

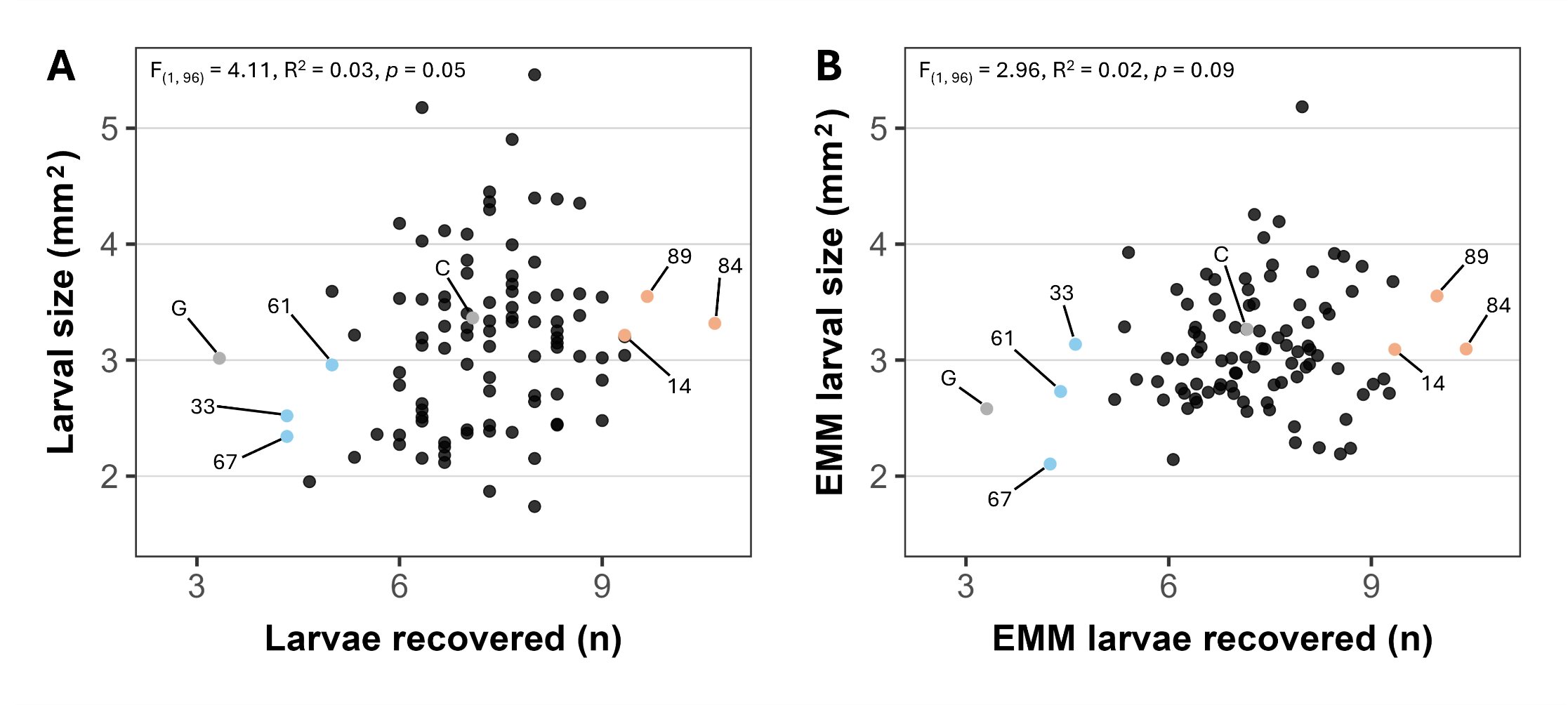
**Figure S15.** Per genotype relationships between cabbage stem flea beetle (*Psylliodes chrysocephala*) larval survival and larval size for all 98 Brassicaceae genotypes (97 *Brassica napus* and a single *Sinapis alba*; **Table S1**) from the Brassicaceae population larval antibiosis screen. **(A)** Raw mean larval size by raw mean larval survival across all 98 Brassicaceae genotypes. **(B)** Estimated marginal mean (EMM) larval size by EMM larval survival across all 98 Brassicaceae genotypes. For both plots, each point represents one of the 98 genotypes included in the diversity screen, with resistant (blue points), susceptible (peach points), and control (grey points) genotypes individually labelled (G = *S. alba* Elsoms G1; C = *B. napus* KWS Campus).

**Figure S16.** Cabbage stem flea beetle (*Psylliodes chrysocephala*) mean adult emergence time (in weeks) across the six selected *Brassica napus* genotypes used for the adult emergence experiment. Genotypes are ordered from lowest to highest mean. Blue and peach points represent genotypes from the *B. napus* population screen that were initially identified as resistant and susceptible, respectively (**Figure 2**). Large points with whiskers represent the mean standard error, jittered points to the left represent the raw data, and letters represent non-significant differences (*p* > 0.05) between groups.
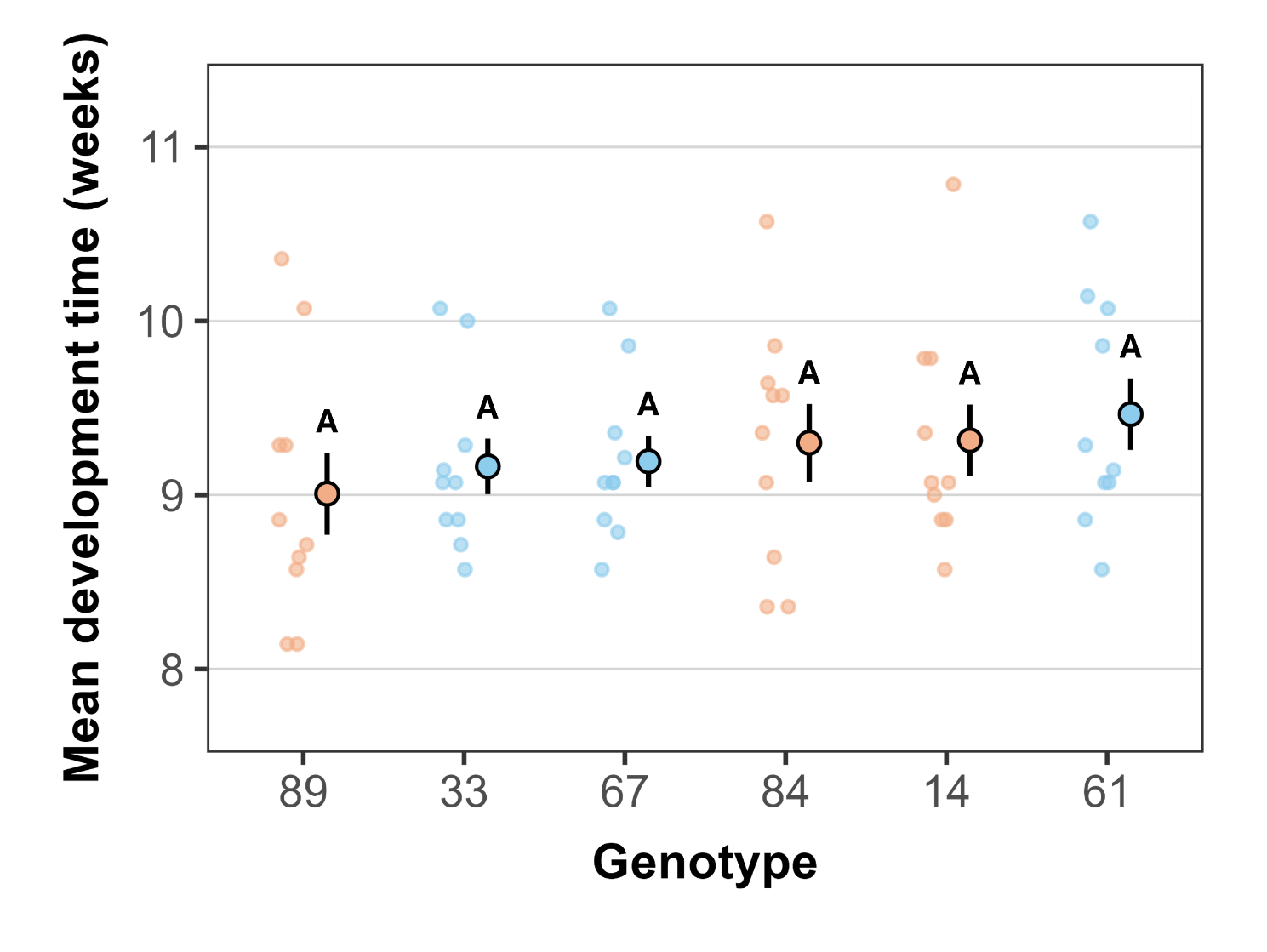
